# Polygenic polymorphisms cooperate to drive expansion of HCMV adaptive NK cell and NK-like T cell populations

**DOI:** 10.64898/2026.11.02.763083

**Authors:** Jean-Benoît Le Luduec, Theodota Kontopoulos, M. Kazim Panjwani, Hongtao Liu, Rosa Sottile, Colleen M. Lau, Katharine C. Hsu

**Author notes:** Corresponding author: Dr. Jean-Benoit Le Luduec Zuckerman 745-D, Memorial Sloan Kettering Cancer Center 408 East 69th street, New York, NY 10021.

## Abstract

Human cytomegalovirus (HCMV) infection drives the expansion of several innate and innate-like lymphocyte populations. Characterized by high cell surface expression of NKG2C, these populations include the “adaptive” CD56^+^ FcεRIγ-NK cell, an NK-like TCRαβ^+^CD8^+^ T cell population and a Vγ9/Vδ1^+^ T cells. Beyond NKG2C expression, both the adaptive NK cell and the NK-like T cell population exhibit enhanced cytokine response to antibody-dependent cellular cytotoxicity (ADCC). Expansion of each population is only observed in 30-40% of HCMV^+^ individuals and is independent of expansion of each other. In this study, we evaluated 200 healthy donors and identified several immunogenetic factors that influence NKG2C expression across the three lymphocyte populations. *NKG2C* (*KLRC2*) copy number and allelic polymorphism was associated with NKG2C^+^ NK cell frequency and surface expression density in HCMV^-^ but not in HCMV^+^ individuals. Investigating other genetic loci encoding proteins related to adaptive NK function, we found that CD16a and HLA-E alleles act cumulatively with *NKG2C* copy number to drive NKG2C expression on lymphocytes among HCMV^+^ individuals. Polymorphism of the NKG2C ligand HLA-E influences strength of NK cell activation via NKG2C, where NK responsiveness was lower in individuals with the HLA-E*01:03 genotype, while CD16a polymorphism influenced strength of ADCC response among NKG2C^+^ TCRαβ^+^CD8^+^ T cells. Based on these observations, we conclude that genetic polymorphisms in *NKG2C*, *CD16a* and *HLA-E* shape the NKG2C^+^ repertoire, influence NKG2C-mediated NK function and dictate capacity of ADCC in response to HCMV infection. Immunogenetic polymorphism among NKG2C-related genes therefore tunes innate immunity to HCMV infection providing additional protection beyond adaptive immunity.

**Visual abstract:** Figure for visual abstract: Sequential activation during primo HCMV infection

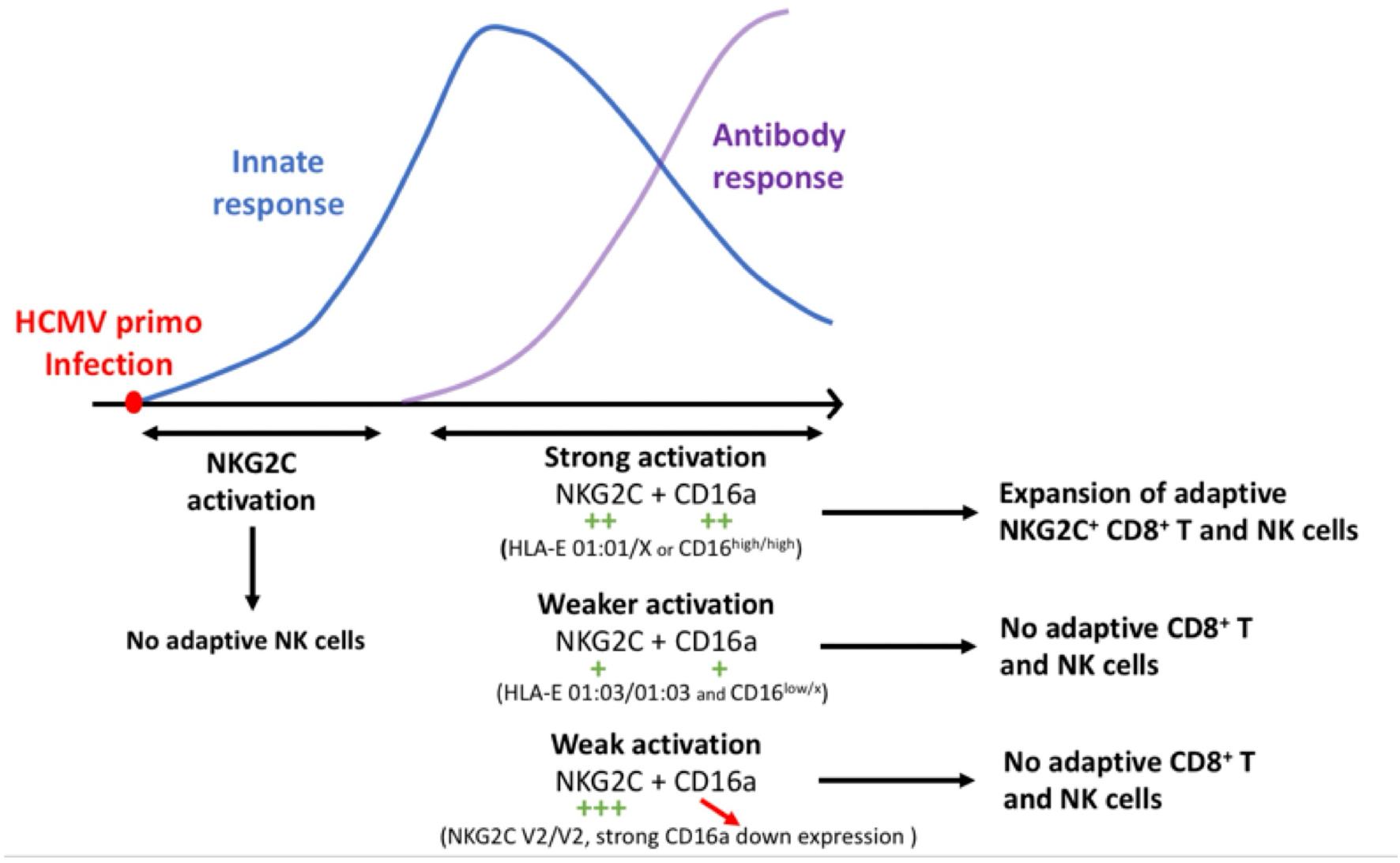

## INTRODUCTION

HCMV is a herpesvirus that causes a persistent lifelong infection and is found with a high worldwide prevalence ranging from 60 to 90% ^1,2^. In healthy individuals, human cytomegalovirus (HCMV) infection is easily controlled by the immune system, driving the expansion of adaptive lymphocyte populations among NK cells and T cells ^3,4^. In immunodeficient or immunosuppressed individuals, initial infection with HCMV or reactivation are of life-threatening concern. The virus causes significant morbidity and mortality in newborn infants infected with HCMV and in allograft recipients ^5^.

HCMV infection induces the development of so-called “adaptive NK cells”, characterized by the co-expression of cellular markers including NKG2C, CD57, and self-HLA specific KIR; the absence of the signaling adaptor FcεRIγ ^6,7^; and a decrease of the transcription factor PLZF, resulting in epigenetic modifications ^8,9^. Functionally, these cells exhibit enhanced cytokine response to antibody-dependent cellular cytotoxicity (ADCC) when compared to non-adaptive NK cells ^6,7^, in addition to heightened cytotoxicity against HCMV-infected fibroblasts ^10,11^. Mature NK cells are universally CD16^+^, and the adaptive FcεR1γ-deficient NK cells have been shown to expand preferentially in an antibody-dependent manner ^8^. Further highlighting the importance of CD16 in shaping the NK cell repertoire and controlling heterologous viral infection, individuals with complete CD16a deficiency exhibit a drastically reduced HCMV-associated adaptive NK population and persistent infection with the Epstein-Barr virus ^12^.

Only 30-40% of HCMV^+^ individuals exhibit an expanded NKG2C^+^ NK cell population ^13,14^. While a study has proposed that recognition of strain-specific UL40 HCMV peptides dictates the heterogeneity of adaptive NKG2C^+^ NK cell populations among HCMV-seropositive individuals ^15^, no host-specific markers have yet been found to predict the likelihood of developing this NK population. Interestingly, a recent study on a cohort of HCMV-seropositive (HCMV^+^) kidney transplant recipients showed a protective effect of preexisting adaptive NKG2C^+^ NK cells on the incidence of posttransplant symptomatic HCMV infection ^16^, reinforcing the importance of understanding the mechanism inducing this cell population.

NKG2C heterodimerizes with CD94, forming an activating receptor that binds the non-classical HLA molecule HLA-E ^17,18^. *KLRC2,* the gene encoding NKG2C, is found on chromosome 12 and is commonly deleted in the general population. Thus, individuals can possess zero, one, or two copies of the gene ^19^, and absence of the gene in allogeneic hematopoietic cell donors has been associated with lower control of HCMV reactivation in hematopoietic cell transplantation ^20^. NKG2C expression on NK cells is not pathognomonic for prior HCMV exposure, as a small NKG2C^dim^ NK population can be observed in HCMV-seronegative (HCMV^-^) healthy donors. In both HCMV^-^ and HCMV^+^ individuals, *NKG2C* copy number influences the frequency and the density of NKG2C expression on the NK cell surface ^21,22^. In addition, NKG2C allelic variation affects the frequency of NKG2C-expressing NK cells in HCMV-seropositive individuals.^23^ Recent studies have shown that HCMV infection not only induces the expansion of an NKG2C^+^ population in the NK cell compartment but is also associated with NKG2C expression in TCRαβ^+^CD8^+^T (CD8^+^ T) cells ^24,25^ and Vγ9/Vδ1^+^ T (Vδ1^+^ T) cells ^26^. We have previously demonstrated that HCMV-associated NKG2C^+^CD8^+^ T cells express several NK cell markers, including CD56 and KIR, and similar to NK cells, exhibit innate functional activity ^24^. In addition to protection from HCMV infection, the NK-like NKG2C^+^CD8^+^ T cells exhibit increased control of mycobacterial infection ^27^. For γδ T cells, an increase in the proportion of effector NKG2C^+^Vδ1^+^ T cells have been observed in heart-lung transplant recipients with HCMV reactivation ^26^.

In the present study, we investigate select single nucleotide polymorphisms (SNPs) in genes encoding receptors or ligands including NKG2C, CD16a and HLA-E, which have been shown to have functional relevance in HCMV-driven adaptive lymphocytes. We demonstrate that these SNPs associate with functional differences that influence the expansion of adaptive NKG2C^+^ NK cells, CD8^+^ T cells, and Vδ1^+^ T cells.

## RESULTS

### NKG2C expression is elevated in multiple lymphocyte populations after HCMV infection but varies among healthy donors

We assessed by flow cytometry the association between prior HCMV infection and the frequency of NK cells, TCRαβ⁺ T cells, and TCRγδ⁺ T cells among total lymphocytes in 204 healthy individuals. While we did not observe any frequency variation for total NK cells, TCRαβ^+^ and TCRγδ^+^ T cells (Supplemental Figure 1A), we did observe differences among the T cell subsets. Among TCRαβ^+^ T cells, we found an increased frequency of CD8^+^and CD4^+^CD8^+^ T cells with a commensurate decrease of CD4^+^ T cells in HCMV^+^ individuals (Supplemental Figure 1B). Among TCRγδ^+^ T cells, we observed a higher frequency of Vδ1^+^ T cells and a lower frequency of Vδ2^+^ T cells in HCMV^+^ individuals in comparison to HCMV^-^ individuals (Supplemental Figure 1C). Among NK cells, we observed no differences between CD56^dim^ and CD56^bright^ subpopulations (Supplemental Figure 1D).

We then measured NKG2C expression among these lymphocyte populations, assessing both frequency and cell surface density. Among individuals, the variance of NKG2C^+^ frequency was greater in NK and Vδ1^+^ T cells than in CD8^+^ T cells (Figure 1A). NKG2C surface density, as measured by mean fluorescence intensity (MFI), was variable between individuals in all three populations, with overall lower expression on NK and Vδ1^+^ T cells compared to CD8^+^ T cells (Figure 1A).

**Fig. 1.**
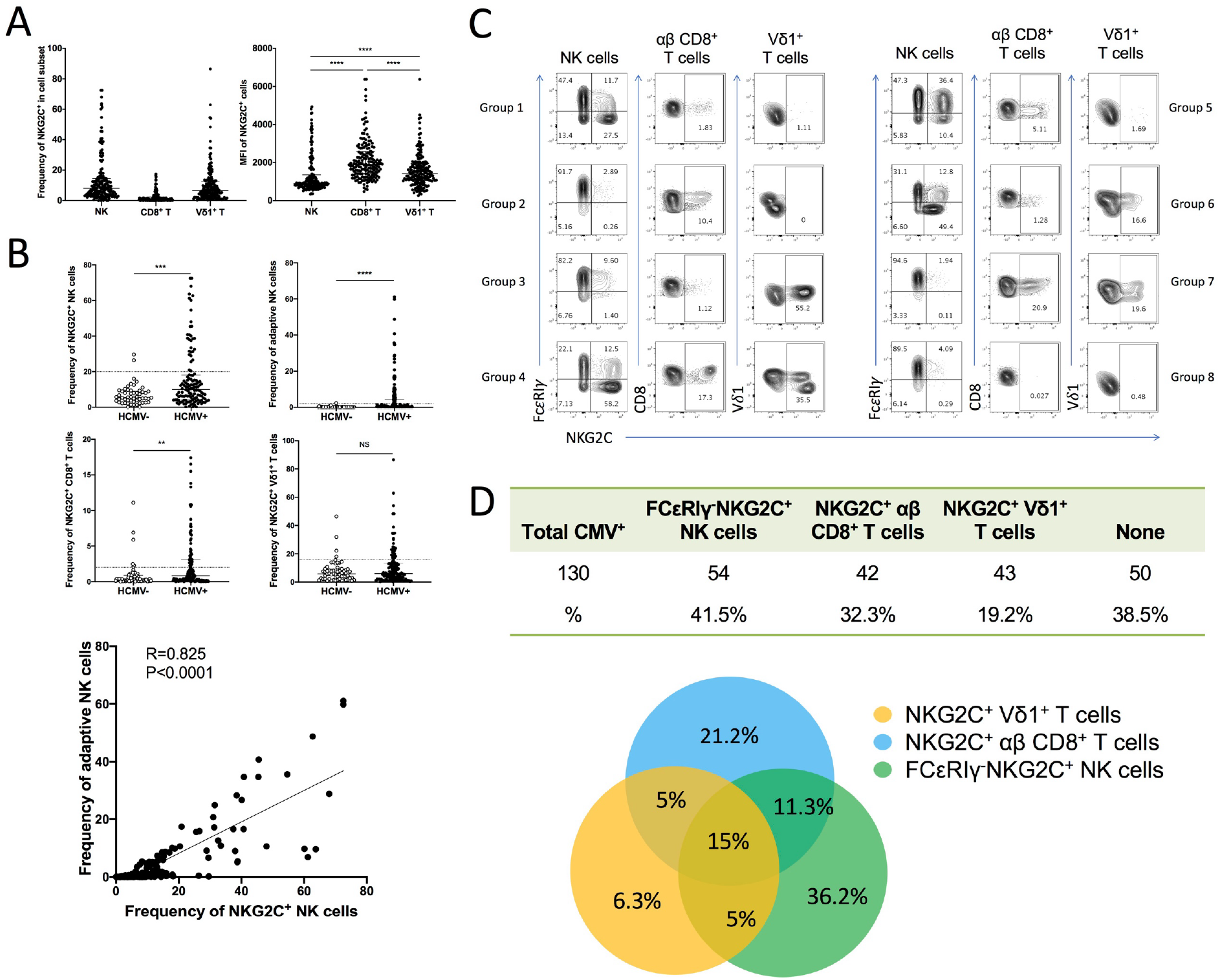
NKG2C expression is elevated in multiple lymphocyte populations after HCMV infection but varies among healthy donors. (A) NKG2C frequency and MFI in NK, CD8^+^ T and Vδ1^+^ T cells. (B) NKG2C frequency stratified by HCMV serostatus in NK, CD8^+^ T and Vδ1^+^ T cells; frequency of adaptive cells stratified by HCMV serostatus in NK cells; correlation between NKG2C^+^ and HCMV adaptive NK cell frequency. (C) NKG2C and FcεRIγ expression in NK, CD8^+^ T, and Vδ1^+^ T cells in 8 representative HCMV^+^ donors. (D) Frequencies of HCMV^+^ individuals with HCMV adaptive induced cells in different lymphocyte populations; distribution of all possible combinations. Mann-Whitney tests were performed and median ± IQRs presented in panels A-F. Correlations were assessed using the Pearson correlation coefficient. Symbols represent individual samples. \**p* < 0.05, \*\**p* < 0.01, \*\*\**p* < 0.001, and \*\*\*\**p* < 0.0001.

Using the frequencies of NKG2C^+^ cell populations observed in HCMV^-^ individuals within each cell population, we established thresholds values in HCMV^+^ individuals indicative of HCMV-induced NKG2C^+^ cell expansion: 20% among NK cells, 2% for adaptive (NKG2C^+^ FcεRIγ^-^) NK cells, 2% among CD8^+^ T cells and 16% among Vδ1^+^ T cells (Figure 1B). We found a strong correlation between the frequency of NKG2C^+^ NK and adaptive NK subsets (Figure 1B). NKG2C frequency was not the only marker associated with HCMV-seropositivity. Frequency of KIR expression, particularly for KIR2DL2/L3/S2, was increased in HCMV^+^ individuals on NK cells in general (Supplemental Figure 1E), TCRαβ^+^ T cells (Supplemental Figure 1F), and TCRγδ^+^ T cells (Supplemental Figure 1G). The increases in KIR expression were only partially since nearly all NKG2C^+^ cells co-express KIR receptors ^13,24^; however, this was not the sole factor, as increased KIR expression was also observed in NKG2C^−^CD8^+^ T cells in HCMV^+^ individuals (Supplemental Figure 1H).

In most cases, HCMV^+^ healthy donors do not develop or expand NKG2C^+^ populations in all three lymphocyte compartments. Among the three cell types, eight theoretical combinations of NKG2C^+^ subpopulations may arise; we observed all eight combinations and calculated their distribution in 130 HCMV^+^ healthy individuals (Figure 1G-H). The majority of HCMV^+^ healthy individuals (61.5%) exhibit one or more HCMV-associated subpopulations, while 38.5% of HCMV^+^ individuals do not have any expanded NKG2C^+^ population. Approximately 15% of individuals expand NKG2C^+^ populations in all three lymphocyte compartments. This study aims to identify the genetic factors that influence NKG2C expression density and frequency among lymphocyte populations.

### *NKG2C* copy number and allelic variation, influence frequency and cell surface expression of NKG2C in lymphocytes populations

Healthy individuals may have zero, one, or two copies of *KLRC2*, the gene encoding NKG2C ^19^, where higher NKG2C copy number is associated with higher frequency and higher cell surface expression of NKG2C in NK cells ^21,23,28^. We find that *NKG2C* copy number variation (CNV) is also associated with frequency and cell surface expression among T cell populations (Figure 2 A-D). In HCMV^-^ individuals, NKG2C expression frequency and cell surface expression on NK cells and CD8^+^ T cells, but not on Vδ1^+^ T cells, were directly related to *NKG2C* CNV (Figure 2A and 2B). In HCMV^+^ individuals, there is an association between NKG2C frequency and density with *NKG2C^+^* NK cells, CD8^+^ T cells and Vδ1^+^ T cells (Figure 2C and 2D). No impact of *NKG2C* CNV was observed on other receptor frequencies in HCMV^+^ lymphocytes (Supplemental Figure 2A, 2B and 2C).

**Fig. 2.**
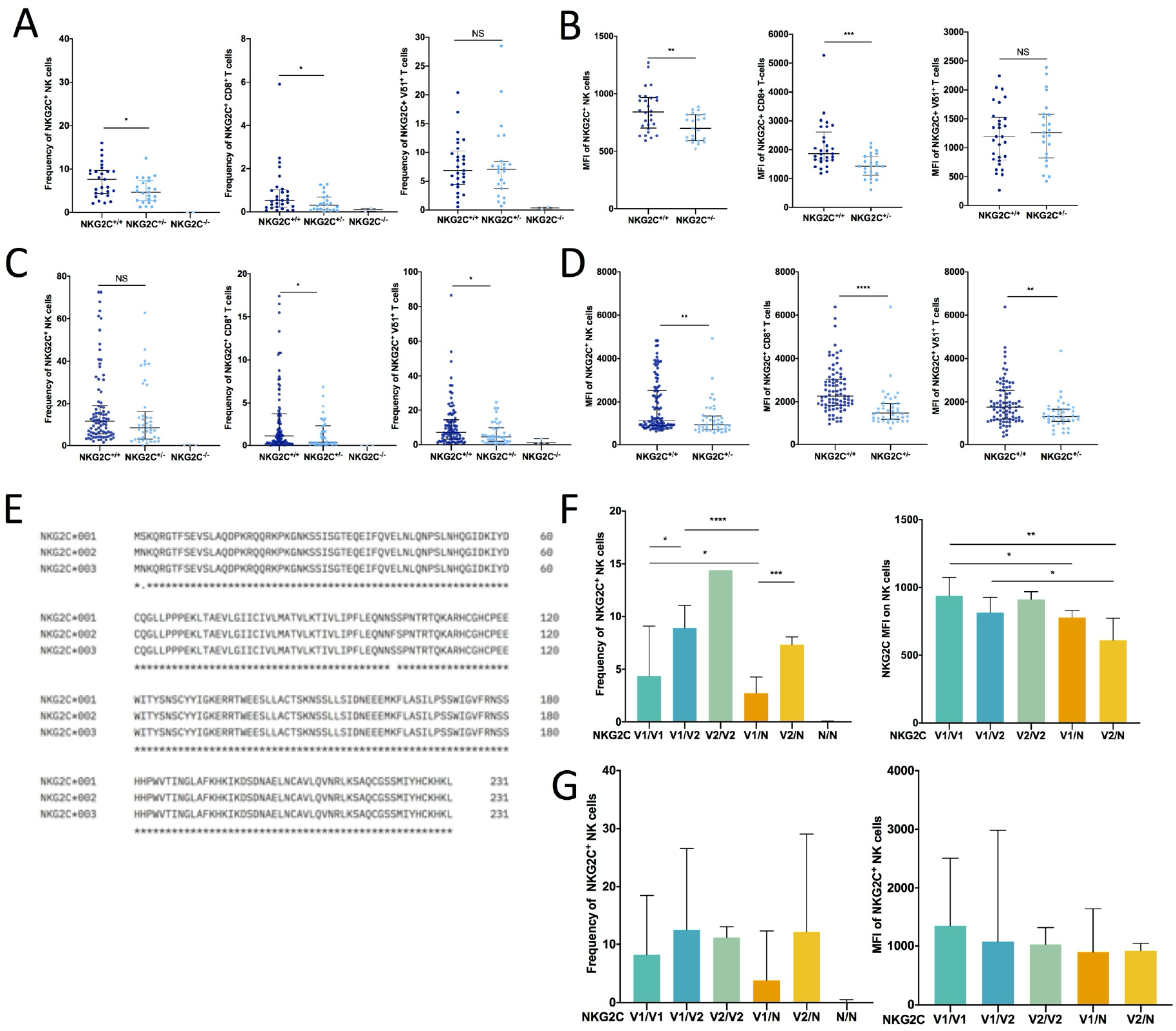
*NKG2C* copy number and allelic variation, influence frequency and cell surface expression of NKG2C in lymphocytes populations. (A) NKG2C frequency stratified by *NKG2C* copy number in NK, CD8^+^ T cells, and Vδ1^+^ T cells in HCMV^-^ individuals. (B) NKG2C MFI stratified by *NKG2C* copy number in NK, CD8^+^ T and Vδ1^+^ T cells in HCMV^-^ individuals. (C) Frequency of NKG2C^+^ CD8^+^ T, and Vδ1^+^ T cells grouped by *NKG2C* copy number in HCMV^+^ individuals. (D) NKG2C MFI stratified by *NKG2C* copy number in NK, CD8^+^ T, and Vδ1^+^ T cells in HCMV^+^ individuals. (E) Protein sequence alignment of the three NKG2C common variants, with changes visible in positions 2 and 102. (F) Frequency and MFI of NKG2C+ NK cells grouped by *NKG2C* copy number and allelic variant in HCMV^-^ individuals. (G) Frequency and MFI of NKG2C^+^ NK cells grouped by *NKG2C* copy number and allelic variant in HCMV^+^ individuals. Mann-Whitney tests were performed and median ± IQRs presented in panels A, B, C, D, F and G. Symbols represent individual samples. \**p* < 0.05, \*\**p* < 0.01, \*\*\**p* < 0.001, and \*\*\*\**p* < 0.0001.

Single nucleotide polymorphism (SNP) at rs1141715 and rs28403159 lead to amino acid changes at positions 2 and 102 of the NKG2C protein sequence, producing three unique NKG2C protein variants ^29,30^ (Figure 2E). Allele typing of the healthy cohort with the 2 SNPs segregated individuals almost identically into variant group 1 (V1) exhibiting allele *NKG2C*01* and variant group 2 (V2) exhibiting allele *NKG2C*02*. Because only one individual was detected with the relatively rare variant *NKG2C*03* ^29,30^, we did not integrate *NKG2C*\**03* in our study as a separate comparison group. Instead, individuals with *NKG2C*03* were included in variant group 1. We evaluated whether allelic variant could further influence NKG2C frequency and MFI beyond NKG2C CNV. In HCMV^-^ individuals, *NKG2C* alleles and gene copy number had a significant combined impact on the frequency of NKG2C^+^ cells, where the V2 variant was associated with increased frequency. CNV and allelic variant did not influence cell surface expression, however (Figure 2F). The same association with frequency was not observed in HCMV^+^ individuals (Figure 2G), nor was there any evident impact of CNV and allele on NKG2C^+^ frequency and surface density on CD8^+^ T cells or Vδ1^+^ T cells regardless of HCMV serostatus (Supplemental Figure 2D-G).

We examined the distribution of *NKG2C* CNV and variants by ethnicity and found an increased frequency of *NKG2C^Neg/Neg^* and variant V2 in self-identified Black/African American donors in comparison to other ethnic identifications (Supplemental Figure 2H).

Overall, we find that *NKG2C* CNV is associated with frequency and cell surface expression of NKG2C in both NK and CD8^+^ T cells in both HCMV^+^ and HCMV^-^ donors, and with frequency and cell surface expression in Vδ1^+^ T cells from HCMV^+^ donors. In contrast, NKG2C allelic polymorphism has limited effect on NKG2C expression, associating only in HCMV^-^ donors with frequency of NKG2C^+^ NK cells. These data suggest that other factors must influence NKG2C expression among lymphocytes.

### CD16a and HLA-E polymorphisms are associated with NKG2C expression on lymphocytes in HCMV^+^ individuals

A recent study highlighted the role of CD16a in the development of adaptive NK, showing that individuals with a complete CD16a deficiency exhibit a drastic reduction in adaptive NK cells frequency ^12^. Furthermore, FcεRIγ-negative NK cells, a commonly used phenotype marker of adaptive NK cells, have been shown to expand preferentially in an antibody-dependent manner ^8^ consistent with their enhanced responsiveness to CD16-mediated activation compared with FcεRIγ-positive NK cells ^7,31^. We therefore hypothesized that an *FCGR3A* polymorphism known to influence CD16a function and phenotype may play a role in the development of adaptive NK cells and other NKG2C^+^ lymphocyte populations. Dimorphism at rs396991 leads to a F158V substitution, with homozygosity for 158V resulting in higher expression and affinity of the CD16a receptor ^32,33^. We examined NKG2C frequency among NK cells and T cell populations in HCMV^+^ donors, segregated by *FCGR3A* polymorphism (Figure 3A, 3B and 3C). Individuals homozygous for the high affinity variant (CD16a V/V) showed a significantly increased frequency of NKG2C^+^ cells in their CD8^+^, TCRαβ^+DP^ and TCRαβ^+DN^ T cells compartment compared to individuals with a low affinity variant. Furthermore, homozygosity for 158V was also associated with an increased frequency of T cells expressing KIR2D (KIR2DL2/L3/S2) receptors, indicating that the effect of *FCGR3A* polymorphism extends beyond NKG2C expression. CD16a polymorphism alone was not significantly associated with NKG2C frequency in NK cells and Vδ1^+^ T cells (Figure 3A and 3B). We considered the variant effects of the NKG2C ligand, HLA-E, on NKG2C expression. The two common allelic variants of HLA-E, 01:01 and 01:03, have lower and higher expression, respectively ^33–36^. We examined NKG2C frequency, stratifying by HLA-E allele (Figure 3D, 3E, and 3F) and observed significantly lower NKG2C frequency among NK cells and CD8^+^ T cells from individuals homozygous for HLA-E*01:03.

**Fig. 3.**
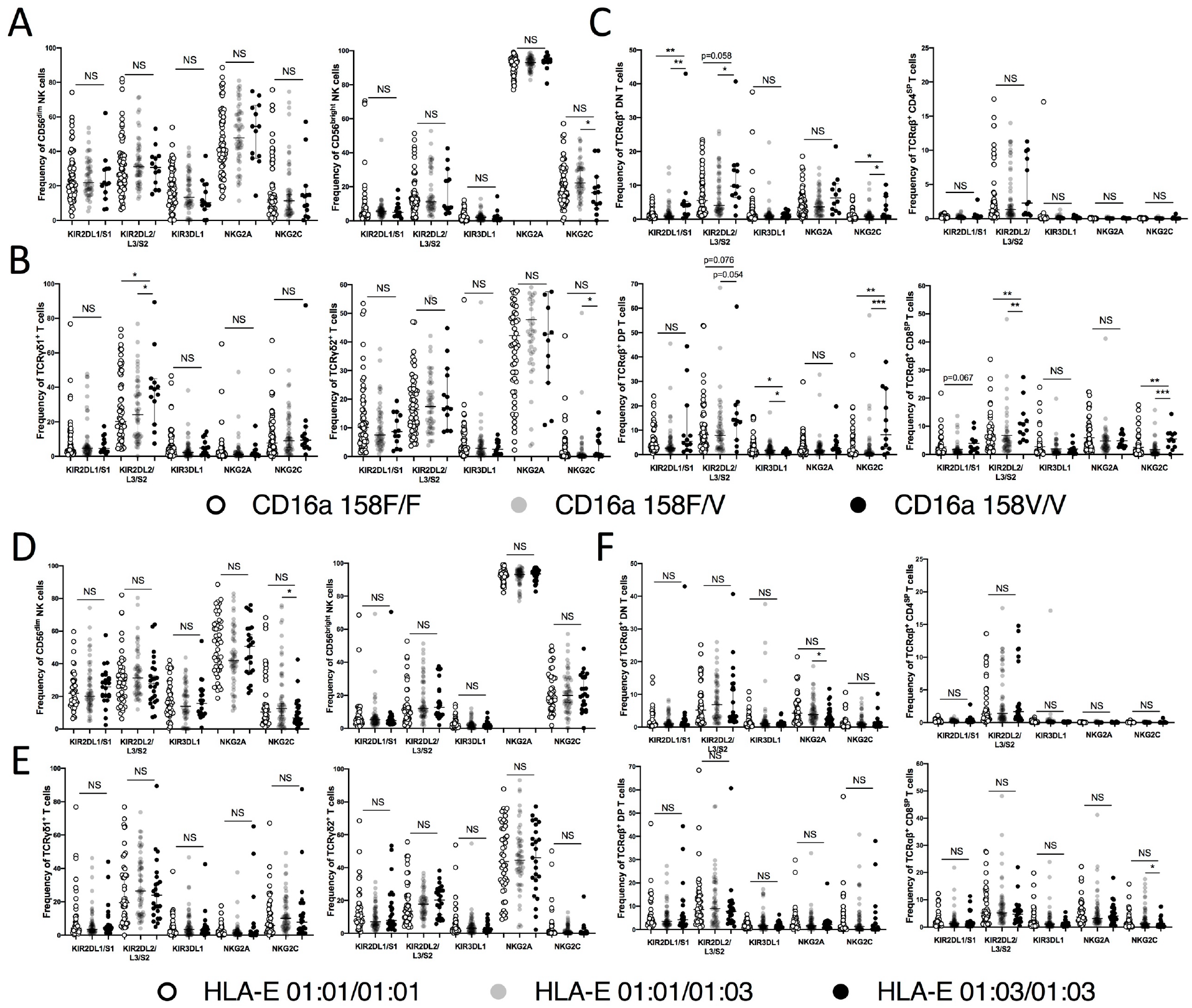
CD16a and HLA-E polymorphisms are associated with NKG2C expression on lymphocytes in HCMV^+^ individuals. NK cells markers expression frequencies stratified by CD16a variants (rs396991 T>G) in HCMV^+^ individuals: (A) in NK cells subsets, (B) in TCRγδ^+^ T cell subsets, (C) in TCRαβ^+^ T cells subsets. NK cells markers expression frequencies stratified by HLA-E variants (rs1264457 G>A) in HCMV^+^ individuals: (D) in NK cells subsets, (E) in TCRγδ^+^ T cell subsets, (F) in TCRαβ^+^ T cells subsets. Mann-Whitney tests were performed and median ± IQRs presented in panels A-F. Symbols represent individual samples. \**p* < 0.05, \*\**p* < 0.01, \*\*\**p* < 0.001, and \*\*\*\**p* < 0.0001.

Due to their potential impact on KIR, HLA-E or NKG2C expression, other immune gene polymorphisms were evaluated but were not found to be associated with NKG2C^+^ lymphocyte expansion. These included polymorphic sites in the HLA-B leader peptide, CD32, CD94, CD2, CD58, HLA-C epitope and NKG2D (Supplemental Figure 3A-G). Overall, we find that individuals homozygous for the CD16a high affinity variant had a significantly increased frequency of NKG2C^+^ and KIR2D^+^ cells among T cell populations, and individuals homozygous for the high-expressing HLA-E variant had a significantly lower frequency of NKG2C^+^ NK and CD8^+^ T cells.

### CD16a, HLA-E and NKG2C polymorphisms cumulatively associate with NKG2C expression on NK and CD8^+^ T cells in HCMV^+^ individuals

We then performed a combined analysis of relevant SNPs to explore their joint association with NKG2C^+^ cell expansion in HCMV^+^ individuals. Polymorphisms of HLA-E and CD16a had a combined impact on NKG2C^+^ cell frequency among NK cells and CD8^+^ T cells but not on Vδ1^+^ T cells (Figure 4A). HCMV^+^ individuals with one or more copies of HLA-E*01:01 and those with two copies of CD16a 158V had a higher frequency of NKG2C^+^ NK cells and CD8^+^ T cells in comparison to the HCMV^-^ group. In individuals lacking HLA-E*01:01 or CD16a 158V, no significant differences were found between HCMV^+^ and HCMV^-^ individuals. This observation held true independent of *NKG2C* copy number (Supplemental Figure 4A). Similar results were obtained for the adaptive NK cell population (Supplemental Figure 4B). Interestingly, among individuals with at least one copy of HLA-E 01:01 or homozygous for CD16a 158V, those homozygous for the higher-expressing NKG2C variant V2 exhibited lower NKG2C expression in NK cells, CD8^+^ T cells (Figure 4B and Supplemental Figure 4C). These findings suggest that the NKG2C V2 variant may influence NKG2C expression across these lymphocyte populations. We did not observe an impact of the combined polymorphism of HLA-E, and CD16a on NKG2C^+^ Vδ1^+^ T cells (Figure 4A-B).

**Fig. 4.**
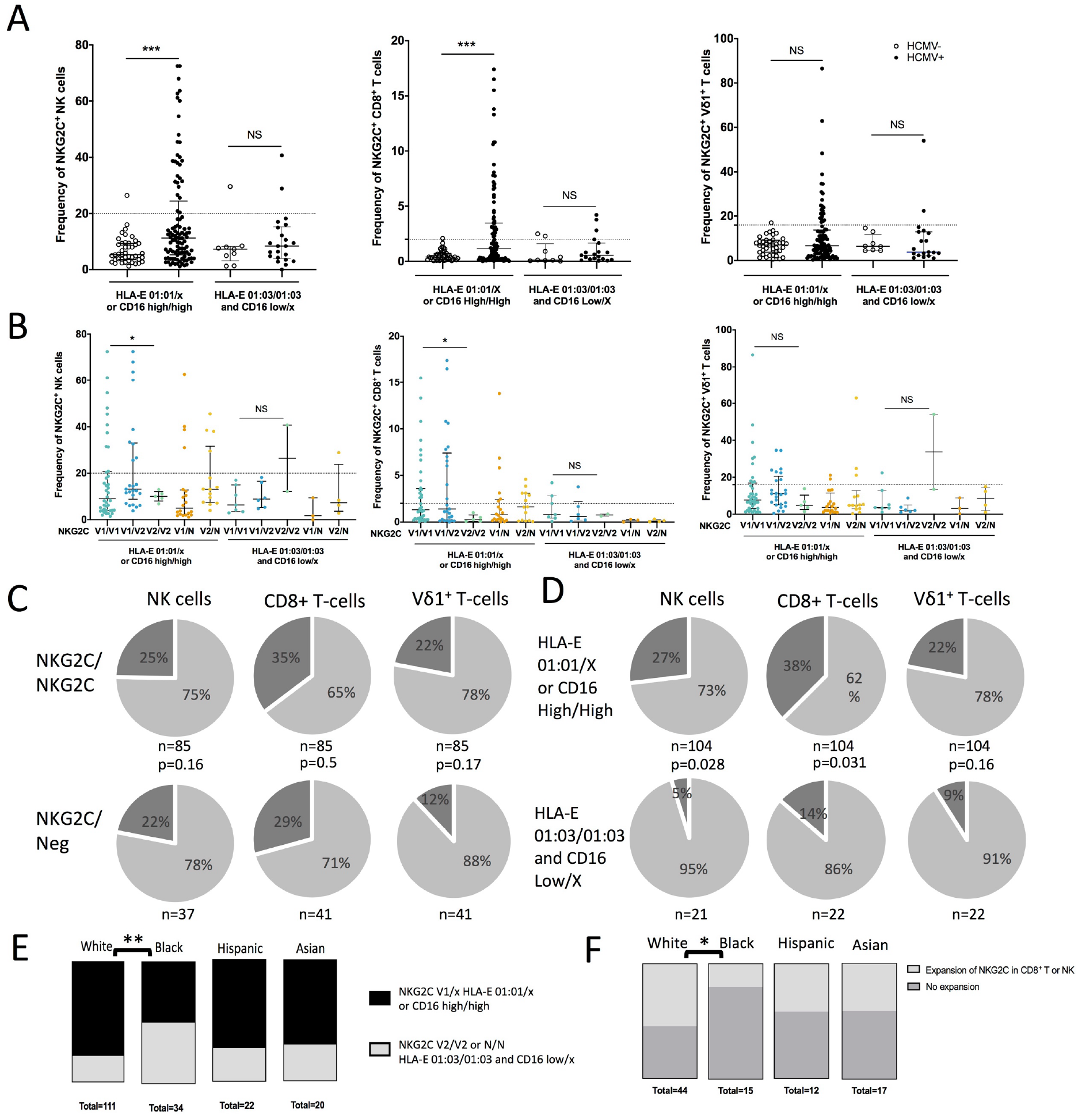
CD16a, HLA-E and NKG2C polymorphisms cumulatively associate with NKG2C expression on NK and CD8^+^ T cells in HCMV^+^ individuals. (A) NKG2C frequency stratified by the combine polymorphism of HLA-E variants (rs1264457 G>A) and CD16a variants (rs396991 T>G) in NK cells, CD8^+^ T cells and Vδ1^+^ T cells from HCMV^-^ and HCMV^+^ individuals. (C) NKG2C frequency stratified by the combine polymorphism of *NKG2C* alleles and copy number, HLA-E variants and CD16a variants in NK cells, CD8^+^ T cells and Vδ1^+^ T cells HCMV^+^ individuals. (C) Distribution of HCMV^+^ individuals having significant NKG2C^+^ cell expansions in NK cells, CD8^+^ T cells or Vδ1^+^ T cells stratified by *NKG2C* copy number. The significance of the NKG2C^+^ was defined by the frequency present in HCMV^-^ individuals. In dark gray individuals having frequency on NKG2C^+^ >20% for NK cells, >2% for CD8^+^ T cells or >16% Vδ1^+^ T cells (F) Distribution of HCMV^+^ individuals having significant NKG2C^+^ cell expansions in NK cells, CD8^+^ T cells or Vδ1^+^ T cells stratified by HLA-E variants and CD16a variants. (D) Distribution of the combine association of *NKG2C* alleles, CD16a and HLA-E alleles among different ethnic groups in our cohort. (E) Distribution of the individuals having expansion NKG2C in NK cells (NKG2C^+^ >20%) or CD8^+^ T cells (NKG2C^+^>2%) among different ethnic groups in our cohort. Individuals with the double deletion of the *NKG2C* gene were excluded from this panel from the analysis. Mann-Whitney tests were performed and median ± IQRs presented in panels A, and B. Comparison of distributions were assessed using Chi-square test in panels C-F. Symbols represent individual samples. \**p* < 0.05, \*\**p* < 0.01, \*\*\**p* < 0.001, and \*\*\*\**p* < 0.0001.

We then investigated the distribution of HCMV^+^ individuals with significant NKG2C^+^ cell expansion among NK, CD8^+^ T, or Vδ1^+^ T cell populations, using the same established thresholds defined in Figure 1B (Figure 4A). No difference in distribution was observed when individuals were stratified by NKG2C copy number (Figure 4C), but the combination of HLA-E and CD16a polymorphisms showed significant associations with NKG2C^+^ cells among NK cells and CD8^+^ T cells (Figure 4D). Based on these results, we can conclude that if *NKG2C* copy number influences the frequency of NKG2C^+^ among NK, CD8^+^ T, and Vδ1^+^ T cells, while HLA-E and CD16a polymorphisms also impact the likelihood of expansion of HCMV-induced NKG2C^+^ cells among NK and CD8^+^ T cells.

We then exanimated the distribution of the combined polymorphisms across ethnic groups. The frequency of these polymorphisms did not differ significantly between HCMV^-^ and HCMV^+^ individuals (Supplemental Figure 4D). However, Black donors had a higher frequency of the genetic markers associated with lower NKG2C expression than White donors (p<0.01) (Figure 4E) and we also observed a significant difference in expansion of NKG2C^+^ NK cell or CD8^+^ T cell between White and Black donors (p<0.05) (Figure 4F).

Overall, we find that CD16a, HLA-E and NKG2C polymorphisms cumulatively associated with NKG2C expression on NK, and CD8^+^ T cells in HCMV^+^ individuals and are differentially distributed between self-identified White and Black donors.

### The expansion of NKG2C^+^ NK cells induced by HCMV associate with CD16a expression and the lack of NKG2A expression

Some receptors, including KIRs, NKG2A, and CD16a, are differentially expressed during NK cell maturation, with CD56^bright^ NK cells generally representing a less mature stage and CD56^dim^ NK cells a more differentiated stage^37^. These receptors can also be co-expressed by NKG2C⁺ NK cells. The majority of CD56^dim^ NK cells have high cell surface expression of CD16a and co-express NKG2A or KIRs, whereas most CD56^bright^ NK cells are CD16a^-^NKG2A^+^KIR^-^ (Figure 5A). Low frequencies of CD56 negative (CD56^neg^) CD16a^+^ NK cells, which express NKG2A or KIRs in a pattern like CD56^dim^ NK cells, have also been observed in healthy individuals ^38,39^.

**Fig. 5.**
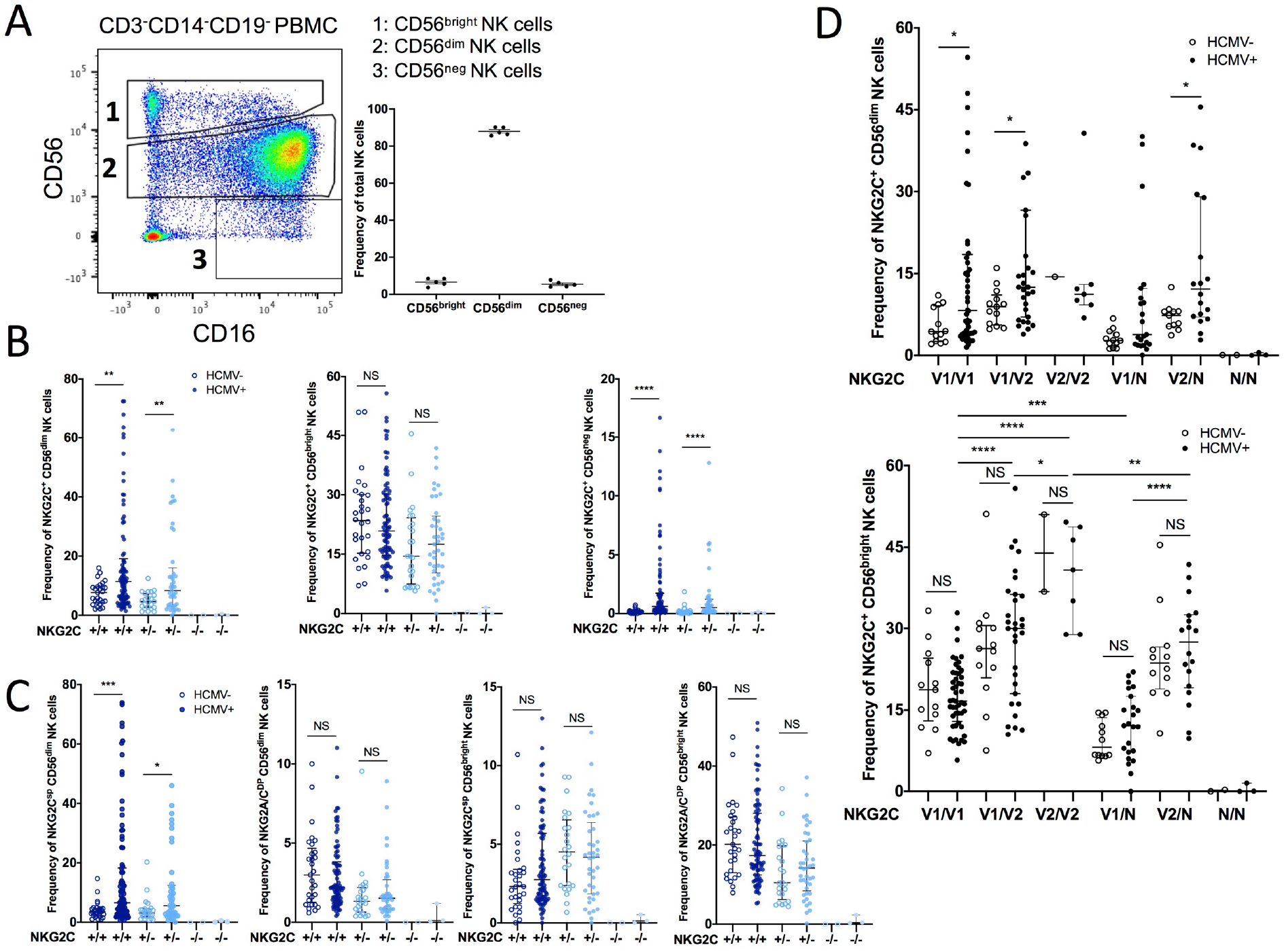
The expansion of NKG2C+ NK cells induced by HCMV correlates with CD16a expression. (A) FACS gating of CD56^bright^, CD56^dim^, and CD56^neg^ NK cells stratified by CD56 and CD16 expression and their relative distribution among total NK cells. (B) NKG2C frequency within the different NK cell compartments stratified by *NKG2C* copy number and HCMV serostatus. (C) NKG2C frequency within CD56^bright^ and CD56^dim^ compartments in NKG2C^SP^ or NKG2A/C^DP^ NK cells, grouped by NKG2C copy number and HCMV serostatus. (D) NKG2C frequency in CD56^dim^ and CD56^bright^ NK cells grouped by NKG2C alleles and copy number and HCMV serostatus. Mann-Whitney tests were performed and median ± IQRs presented in panels B-D. Symbols represent individual samples. \**p* < 0.05, \*\**p* < 0.01, \*\*\**p* < 0.001, and \*\*\*\**p* < 0.0001.

Given the expression of NKG2C across the NK cell populations defined by CD56 expression (bright, dim or neg), we sought to determine whether HCMV serostatus, co-expression of NKG2A or CD16a, and genetic polymorphisms influence NKG2C expression within each of these NK cell populations.

Prior HCMV infection influenced NKG2C expression frequency within different NK cell developmental populations, but not all (Figure 5B). We observed a significant increase in NKG2C frequency among CD56^dim^ and CD56^neg^ NK cells in HCMV^+^ individuals, while there was no difference in NKG2C frequency among CD56^bright^ NK cells between HCMV^-^ and HCMV^+^ individuals. Although NKG2C and NKG2A expression are generally considered mutually exclusive ^40^, a small but measurable population of CD56^dim^ and CD56^bright^ NK cells can co-express both receptors ^41^. There was no difference in frequency of the NKG2A^+^NKG2C^+^ NK populations, CD56 bright or dim, attributed to HCMV infection (Figure 5C). In contrast to CD56dim NK cells, CD56bright NK cells showed no significant difference in NKG2C expression between HCMV-negative and HCMV-positive donors, consistent with their low CD16a expression. The frequency of NKG2A/NKG2C co-expression was similar between HCMV^-^ and HCMV^+^ individuals in both the CD56^dim^ and CD56^bright^ subsets. These finding suggest that the inhibitory signal from NKG2A may override the activating signal from NKG2C and CD16a, potentially limiting NKG2C-driven expansion in subsets that co-express NKG2A.

When analyzing the same data stratified by the *NKG2C* alleles (Figure 5D), we observed that NKG2C allelic polymorphism was associated with the frequency of NKG2C^+^ CD56^bright^ and NKG2C^+^ CD56^dim^ NK cells. Notably, a significant allelic impact was detected only in HCMV^-^ individuals within the CD56^dim^ NK subset. In addition, the frequency of NKG2A/C^DP^ CD56^dim^ NK cells was influenced by *NKG2C* alleles in both HCMV^-^ and HCMV^+^ individuals (Supplemental Figure 5).

Overall, our findings suggest that following HCMV infection, the expansion of NKG2C^+^ NK cell is associated with the co-expression of activating receptors--CD16a and NKG2C--and the absence of NKG2A. In contrast, in the absence of NKG2C^+^ expansion, NKG2C frequency appears to be primarily determined by *NKG2C* alleles and CNV.

### CD16a receptor expression on CD8^+^ T cells correlates with NKG2C and KIR expression and mediates ADCC in individuals carrying the CD16a 158V genotype

Given the association between the CD16a 158V/V polymorphism and NKG2C^+^ CD8^+^ T cell expansion, we investigated CD16a function on these cells. NKG2C^+^ CD8^+^ T cells from HCMV^+^ individuals express the highest level of CD16a compared to NKG2A^+^ or NKG2A/C^DP^ CD8^+^ T cells (Figure 6A). NKG2C^+^ CD8 T cells from individuals homozygous for the high-expression allele CD16a 158V/V exhibiting the highest CD16a surface density and frequency (Figure 6B and Supplemental Figure 6A-D). Furthermore, addition of IL-15 disproportionately induced CD16a expression on 158V/V CD8 T cells from HCMV+ individuals (Figure 6B). In contrast, CD16a expression was low in CD8^+^ T cells of HCMV^-^ individuals and remained unchanged upon addition of IL-15 (Supplemental Figure 6E).

**Fig. 6.**
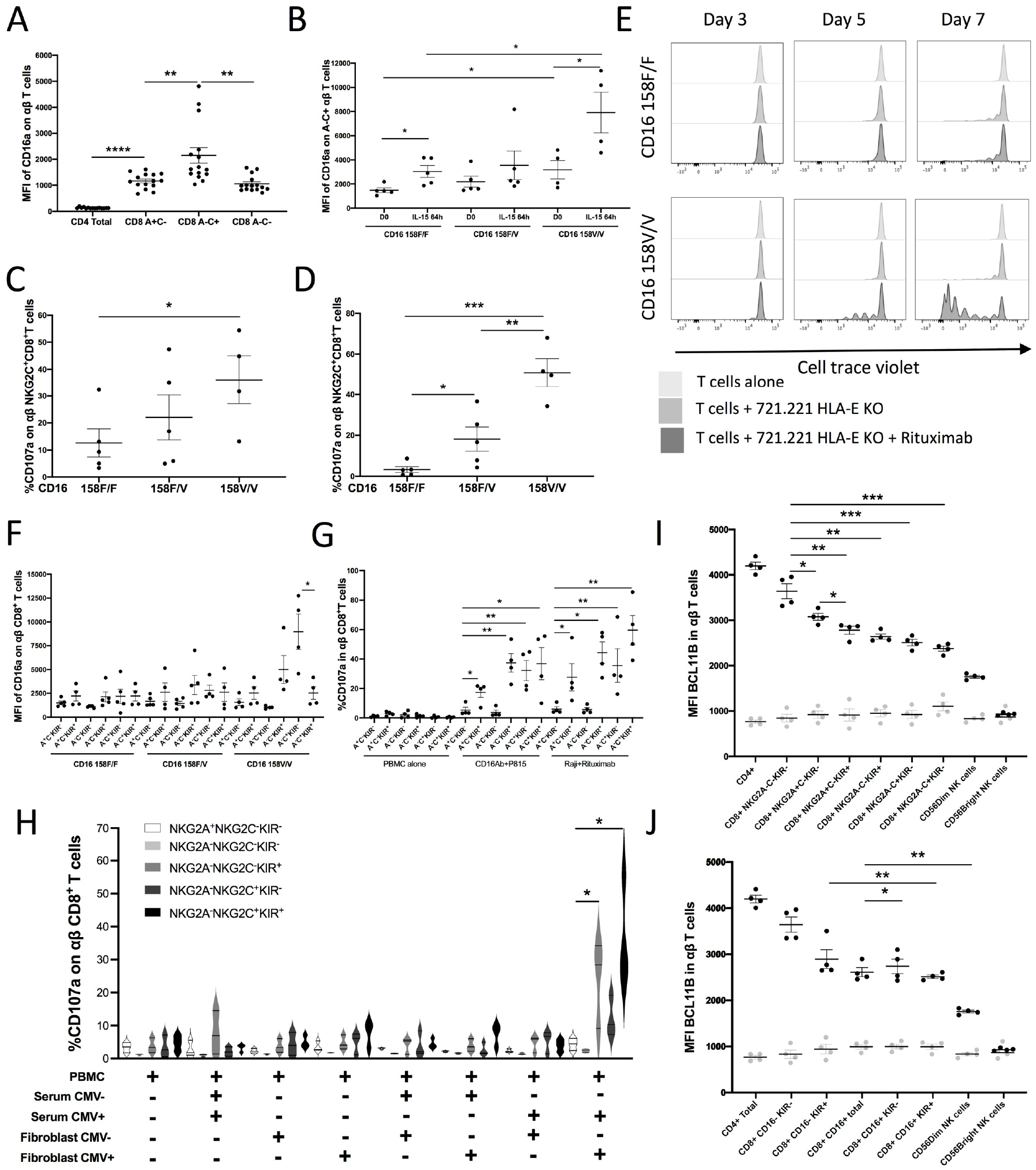
CD16a receptor expression on CD8^+^ T cells correlates with NKG2C and KIR expression and mediates ADCC in individuals carrying the CD16a 158V genotype. (A) CD16a MFI of CD4^+^ or CD8^+^ T cells subpopulation from HCMV^+^ individuals, defined by NKG2A and NKG2C positivity, before IL-15 stimulation. (B) CD16a MFI of NKG2A^-^ NKG2C^+^ CD8^+^ T cells subpopulation from HCMV^+^ individuals with different CD16a polymorphisms, before and after 64 hours of IL-15 stimulation. (C) CD107a response of NKG2A^-^ NKG2C^+^ CD8^+^ T cells subpopulation from HCMV^+^ individuals grouped by CD16a polymorphisms, after 64 hours of IL-15 stimulation, co-cultured with anti-CD16a and P815 cells for 4 hours. (D) CD107a response of NKG2A^-^ NKG2C^+^ CD8^+^ T cells subpopulation from HCMV^+^ individuals grouped by CD16a polymorphisms, after 64 hours of IL-15 stimulation, co-cultured with Raji cells and anti-CD20 Rituximab mAb. (E) CD8^+^ T proliferation assay of negatively selected Pan T cells from peripheral blood from CD16a 158F/F and CD16a 158V/V HCMV^+^ individuals with or without irradiated CD20^+^ 721.221 HLA-E KO in absence or presence of anti-CD20 Rituximab mAb in cRPMI supplemented with IL-15. (F) CD16a MFI of CD8^+^ T cells subpopulation from HCMV^+^ individuals with different CD16a polymorphisms, defined by NKG2A, NKG2C and KIR positivity, after 64 hours of IL-15 stimulation. (F) CD107a response of CD8^+^ T cells subpopulation from HCMV^+^ individuals with different CD16a polymorphisms, defined by NKG2A, NKG2C and KIR positivity, after 64 hours of IL-15 stimulation, co-cultured with indicated conditions. (G) CD107a response of CD8^+^ T cells subpopulation from CD16a 158V/V HCMV^+^ individuals, defined by NKG2A, NKG2C and KIR positivity, after 64 hours of IL-15 stimulation, co-cultured with fibroblasts infected or not by HCMV in presence or not of serum of HCMV^-^ or HCMV^+^ individuals. (I) Bcl11b MFI and isotype control in TCRαβ^+^CD4^+^, CD8^+^ T cells and NK cells subpopulation from CD16a 158V/V HCMV^+^ individuals, defined by KIR, NKG2A and NKG2C positivity (Isotype in gray, Bcl11b in black). (J) Bcl11b MFI and isotype control in TCRαβ^+^ CD4^+^, CD8^+^ T cells and NK cells subpopulation from CD16a 158V/V HCMV^+^ individuals, defined by KIR and CD16a positivity (Isotype in gray, Bcl11b in black). Mann-Whitney tests were performed and median ± IQRs presented in panels A-E and G-K. Symbols represent individual samples. \**p* < 0.05, \*\**p* < 0.01, \*\*\**p* < 0.001, and \*\*\*\**p* < 0.0001.

We then evaluated the impact of CD16a polymorphism on antibody-mediated activation of NKG2C^+^ CD8^+^ T cells. To assess the capacity of response following CD16 engagement, we crosslinked the molecule with anti-CD16a and the murine mastocytoma cell line P815 in a reverse ADCC assay. Activation of CD8^+^ T cells was observed across all CD16a polymorphism groups, with the highest activation seen in cells from CD16 158V/V individuals (Figure 6C and Supplemental Figure 6F). When measuring ADCC with CD20^+^ Raji cells and anti-CD20 Rituximab, however, only NKG2C^+^ CD8^+^ T cells from individuals homozygous for the high expressing CD16a allele 158V exhibited strong ADCC activity (Figure 6D and Supplemental Figure 6G).

We then compared the proliferative capacity of CD16a 158F/F vs CD16a 158V/V T cells following ADCC stimulation from HCMV^+^ individuals by co-culturing the cells with the CD20^+^ cell line 721.221 with or without the anti-CD20 antibody Rituximab. We observed proliferation in CD8^+^ T cells from CD16a 158V/V individuals, but not from CD16a 158F/F individuals (Figure 6E).

As CD16a expression is associated with KIR expression on CD8^+^ T cells ^42^, we conducted additional experiments to determine how KIR co-expression modifies CD16a function, particularly among the NKG2C^+^ population. We segregated CD8^+^ T cells based on expression of NKG2A and NKG2C, as well as KIR2DL1/S1, KIR2DL2/L3/S2, and KIR3DL1. We found that CD16a expression on CD8^+^ T cells was associated not only with KIR expression but also with NKG2C expression. Among CD8^+^ T cells, the NKG2C^+^KIR^−^ subset exhibited the highest levels of CD16a compared to the NKG2C^−^KIR^+^ or NKG2C^+^KIR^+^ subpopulations (Figure 6F). When assessing the function of these cells by crosslinking CD16a or by measuring ADCC activity, we observed that despite having lower CD16a expression, both NKG2C^−^KIR^+^ and NKG2C^+^KIR^+^ subpopulations were more functional than the NKG2C^+^KIR^−^ cells (Figure 6G and supplemental Figures 6H-I). Similar activation patterns were observed when stimulating the cells with HCMV-infected fibroblasts with the serum of HCMV^+^ donors (Figure 6H).

BCL11B, a key transcription factor for T cell lineage, is significantly lower in NK cells and in NKG2C^+^ CD8^+^ T cells ^24^ and other CD8^+^ T cells expressing NK cell associated receptors ^43,44^. We measured the intracellular expression of BCL11B by flow cytometry across different CD8^+^ T cell subpopulations (Figure 6I) and observed a decrease in BCL11B expression that correlated with the presence of NK cell receptors. The more CD8^+^ T cells expressed markers associated with NK cell maturation, such as KIR and NKG2C, the lower their BCL11B expression. Among these markers, CD16a and KIR were most strongly associated with the downregulation of BCL11B in CD8^+^ T cells (Figure 6J).

Altogether, these findings provide insight into the relationship between CD16a polymorphism and the development of the NKG2C^+^ CD8^+^ T cell population in HCMV^+^ individuals. Furthermore, they demonstrate that the acquisition of an NK-like phenotype is linked to the NK-like functional properties of CD8^+^ T cells.

### NKG2C receptor function is influence by both NKG2C and HLA-E allelic variation and acts additively with CD16a to enhance NK cell activation

We have previously demonstrated that NK cells expressing activating KIR2DS1 are tolerized by environmental HLA ligand. We inquired whether the responsiveness of NKG2C^+^ NK cells is similarly impacted by environmental HLA-E.

We developed a functional assay using PBMCs from HCMV^-^ individuals co-cultured with 721.221 cells transduced to express HLA-E and various nonameric HLA class I peptides. CD107a and IFN-γ mobilization were observed in NKG2C^SP^ NK cells only in response to the VMAPRTLFL peptide. In contrast, stimulation with VMAPRTLVL and VMAPRTLIL resulted in baseline levels of activation, comparable to the 721.221 HLA-E knockout cells. Among NKG2C^+^ NK cells responding to VMAPRTLFL, we detected differential activation within NKG2C*01 donors, where HLA-E*01:01 homozygosity individuals had significant higher responsiveness compared to HLA-E*01:03 homozygotes (p<0.05) (Figure 7A). Interestingly, individuals carrying the NKG2C*02 variant and HLA-E 01:01 homozygosity demonstrated even greater activation than those with NKG2C*01 and HLA-E*01:01 (p<0.05) (Figure 7A). To confirm that the differential responsiveness was specific to the HLA-E/NKG2C axis, we evaluated NKG2C^+^ NK cell activation via CD16 cross-linking (Supplemental Figure 6). No significant differences in the CD107a mobilization were observed between HLA-E*01:01 and HLA-E*01:03 homozygote under these conditions. These results indicate that HLA-E allelic variation modulate NKG2C^+^ NK cell responsiveness, and that NKG2C variants differ in their capacity to respond to specific the HLA-E:peptide complex.

**Fig. 7.**
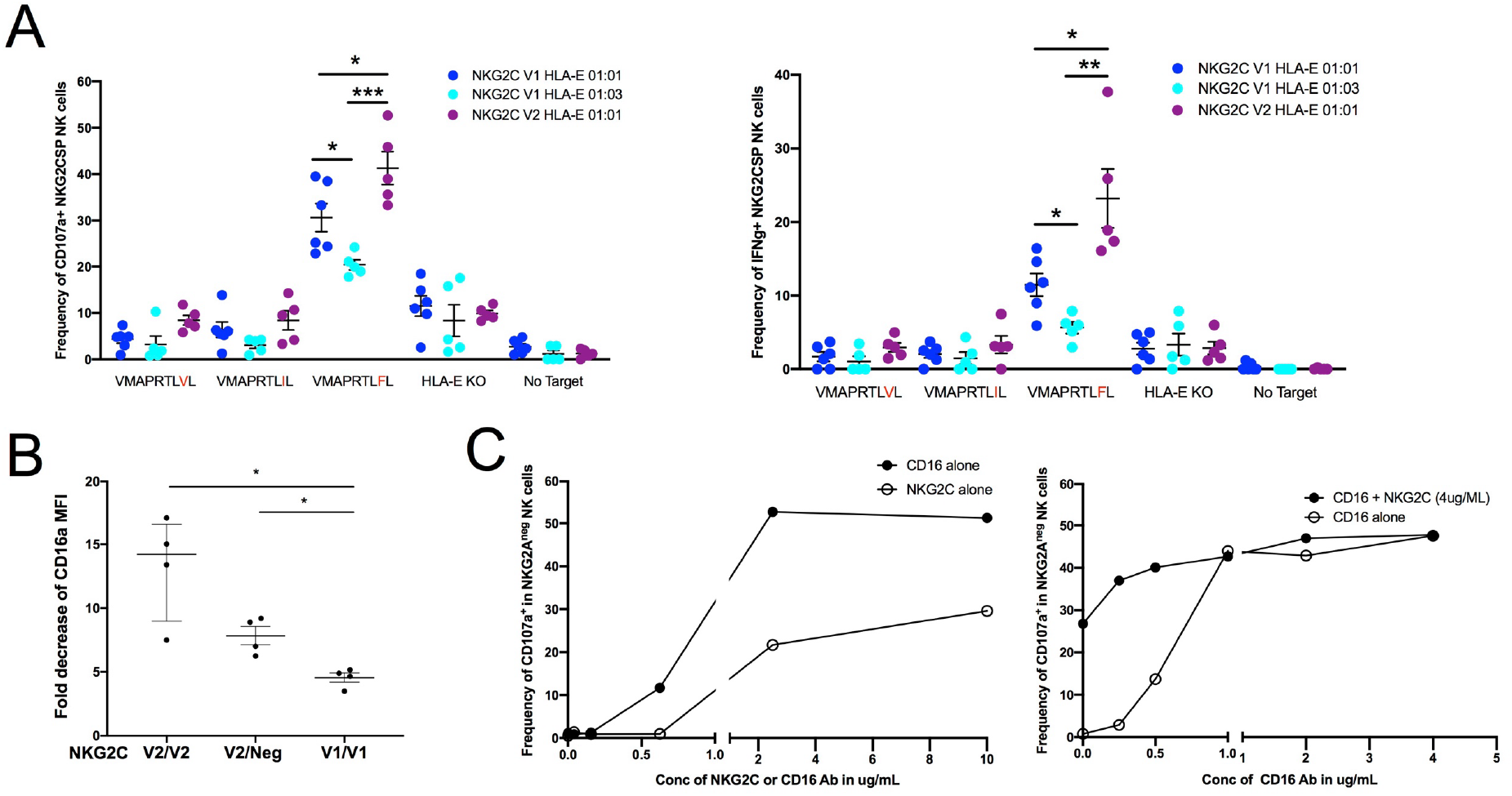
NKG2C receptor function is influence by both NKG2C and HLA-E allelic variation and acts additively with CD16a to enhance NK cell activation. (A) CD107a and IFN-γ response of NKG2C^SP^ NK cells following activation with 721.221 target cells expressing HLA-E and a UL40 peptide, stratified by HLA-E variant (rs1264457 G>A) and *NKG2C* variant and copy number. (B) Fold decrease of CD16a cell surface expression between CD107a^+^ and CD107a^-^ on NKG2C^SP^ NK, from HLA-E*01:01 positive individuals, activated overnight with anti-NKG2C loaded P815 target cells. (C) NKG2A negative (NKG2A^neg^) NK cell CD107a response to plate bound activation assays representative of 3 independent experiments. T-tests were performed and mean ± SEM are presented in panel A. Mann-Whitney tests were performed and median ± IQRs presented in panel B. Symbols represent individual samples. \**p* < 0.05, \*\**p* < 0.01, \*\*\**p* < 0.001, and \*\*\*\**p* < 0.0001.

Using the 721.221 cell line engineered to express HLA-E presenting the VMAPRTLFL peptide, we compared the activation-induced downregulation of surface CD16a in the NKG2C^+^ NK cells from HLA-E*01:01 homozygotes, HCMV^-^ individuals carrying various NKG2C genotype (V2/V2, V2/Neg or V1/V1). CD16a is known to be shed from NK cell surface following activation ^45^. To quantify this effect, we calculated the fold decrease in CD16a expression between activated (CD107a^+^) and non-activated (CD107a^-^) NK cells in overnight in co-culture with 721.221 HLA-E:VMAPRTLFL. We found that NKG2C^+^ NK cells from *NKG2C* V2/V2 individuals exhibited the greatest reduction in CD16a surface expression (Figure 7B) potentially impairing their capacity for ADCC.

Next, we investigated weather co-activation of CD16a and NKG2C receptors has a cumulatively effect on NK cell activation. This experiment was performed using HCMV^+^ individuals with a high frequency of NKG2A^-^NKG2C^+^ NK cells, as plate-bound activation with NKG2C monoclonal antibody (mAb) precludes subsequent detection of NKG2C surface expression. We first tested NK cell activation using plate-bound anti-CD16a and anti-NKG2C mAb across a range of concentrations to determine optimal activation conditions. Subsequently, we assessed CD107a mobilization in response to a fixed dose of anti-NKG2C mAb combined with a serial dilution of anti-CD16a mAb. This co-stimulation resulted in a greater CD107a response compared to NKG2C alone, indicating an additive effect of CD16a and NKG2C co-activation (Figure 7C).

Overall, we find that both NKG2C and HLA-E polymorphism influence NKG2C^+^ NK cell function, and that NKG2C co-expression enhances CD16a-mediated NK cell activation. Altogether, these findings provide insight into the relationship between NKG2C and HLA-E polymorphisms and the development of NKG2C^+^ lymphocytes in HCMV^+^ individuals.

## DISCUSSION

NKG2C can be found on NK cells, as well as on CD8^+^ T cells and Vδ1^+^ T cells ^46–49^, but no study has conducted a systematic analysis examining expansion of NKG2C^+^ cells in these three lymphocyte populations. We report that following HCMV infection, most healthy individuals develop a substantial HCMV-induced NKG2C^+^ population in one of these lymphocyte compartments and more than one NKG2C^+^ compartment may expand simultaneously within the same individual. The HCMV adaptive NK cell population is reported to be stable over time in healthy donors ^13,31^, but it is possible that contraction of the population after infection may explain the 38.5% of HCMV^+^ individuals without an NKG2C^+^ population detectable using current flow cytometrically based methodologies.

*KLRC2* rs1141715 A>G SNP combined with CNV predicts NKG2C^+^ cell frequency in both CD56^dim^ and CD56^bright^ NK cells from HCMV^-^ individuals, as well as in CD56^bright^ NK cells from HCMV^+^ individuals. The limited expansion of NKG2C^+^ cells within the CD56^bright^ compartment may reflect the distinct functional and developmental characteristics of these cells, including lower CD16a expression and frequent NKG2A co-expression. Alternatively, NKG2C^+^ CD56^bright^ cells may represent an intermediate differentiation stage that subsequently transitions to a CD56^dim^ phenotype.

We observed a differential distribution of *NKG2C* alleles across ethnic groups. Both the *NKG2C* gene deletion and the V2 variant were more frequent among the self-identified Black individuals compared to those who self-identified White, Asian, and Latino. Although our cohort size is limited, these observations are consistent with data reported in larger datasets available through the NCBI SNP database (https://www.ncbi.nlm.nih.gov/snp/). In these datasets, the allele rs1141715 G was present at an average of 41.67% in a cohort of 9,912 European individuals and at 83.27% in a cohort of 5,928 Black/African American individuals.

We observed that CD16a polymorphism impacts the frequency of HCMV-adaptive NK cells and NKG2C^+^ CD8^+^ T cells, with a notably increased frequency of NKG2C^+^ NK cells in individuals homozygous for the CD16a 158V allele, the high-affinity variant of this Fc receptor. Given that adaptive NK cells are potent ADCC effectors and expand preferentially in response to antibody stimulation ^8^ and that individuals with a complete CD16a deficiency exhibit a markedly reduced HCMV-adaptive NK cell population ^12^, we hypothesized that CD16a affinity may influence the development of adaptive lymphocyte subsets. Within this framework, the NKG2C receptor may contribute to an alternative activation pathway or as a co-stimulatory signal. Notably, we did not observe NKG2C^+^ expansion in CD16a^-^ CD56^bright^ NK cells, even in individuals with a robust HCMV-induced expansion of NKG2C^+^ CD16a^+^ CD56^dim^ NK cells.

It has been reported that individuals with a homozygous deletion of *NKG2C* can develop a CD57^+^ self-KIR^+^ adaptive NK cell population lacking NKG2C expression and yet remain free of HCMV complications when otherwise healthy ^14^. However, *NKG2C* deletion has been associated with an increased risk of HCMV reactivation and HCMV disease in immunocompromised individuals following HSCT ^20,50^ and has also been studied in the context of HIV infection and solid organ transplantation ^51,52^.

We were particularly interested to find an impact of CD16a polymorphism on expanded NKG2C^+^ CD8^+^ T cells, as few individuals in our healthy cohort displayed baseline CD16a expression on these cells. Although CD16a expression on T cells was first reported decades ago ^53,54^, it remains best characterized in TCRγδ T cells ^55–57^, where its expression is more consistently observed. Recent studies have identified NK-like CD8^+^ T cells expressing CD16a in several disease contexts, including chronic hepatitis C ^58^ HIV ^59^, cancer ^60^, chronic graft-versus host-disease (GVHD) in kidney transplant recipients ^61^, severe COVID-19 ^62^ and in utero HCMV infection ^43^. In these settings, CD16a was reported to be functionally active and capable of mediating ADCC in TCRαβ^+^CD8 T cells ^43,59,62^.

In our study, we observed increased *FCGR3A* transcription in HCMV-induced NKG2C^+^ CD8^+^ T cells ^24^. As none of the donors used in our study were likely experiencing active HCMV infection, we hypothesize that CD16a expression on CD8^+^ T cells may be a response to chronic inflammation. Supporting this, previous studies reported that CD16a^+^ CD8^+^ T cells respond to IL-12/IL-18 stimulation ^60^ and that *in vitro* stimulation of PBMC with EBV-infected B cells and IL-2 can induce CD16a expression of on CD8^+^ T cells ^63^. Moreover, a recent study showed that IL-15 enhances CD16a expression on KIR^+^ CD8^+^ T cells, augmenting their ADCC response against Rituximab-coated 721.221 cells ^42^.

We were able to reproduce this finding, demonstrating robust ADCC in NKG2C^+^ CD8^+^ T cells, but only in individuals homozygous for the high-affinity CD16a 158V variant. The ADCC response was confined to the CD16a^+^ CD8^+^ T cell subset, although the magnitude of the response did not correlate directly with CD16a surface levels. Interestingly, stronger ADCC responses were observed in KIR-expressing CD8^+^ T cells. This association suggests that KIR expression may identify a more differentiated or functionally specialized CD8^+^ T-cell population with enhanced ADCC capacity. To our knowledge, this is the first report linking functional enhancement in TCRαβ^+^ T cells to the expression of inhibitory KIR.

We report reduced NKG2C-mediated function in individuals homozygous for HLA-E*01:03 allele, as compared to those homozygous for HLA-E*01:01, even though 01:03 is associated with higher expression of HLA-E. This seemingly paradoxical finding may be explained by a mechanism of tolerization, similar to what has been described for KIR2DS1, another activating NK receptor that tolerizes its ligand (HLA-C2) in a dose dependent manner ^64,65^. This tolerization mechanism, which can be considered as “reverse education”, is conceptually the inverse of a classical NK education occurring via inhibitory KIRs or NKG2A receptors ^34,66–70^. Reverse education may function as a safeguard to prevent chronic activation of NK cells, thereby reducing the risk of autoimmunity. This concept could help explain our finding that HLA-E*01:03 homozygosity is associated with lower frequency of HCMV-adaptive NK and CD8^+^ T cells. In this scenario, the damped responsiveness of NKG2C^+^ NK cells against HLA-E may limit the development NKG2C^+^ HCMV-induced lymphocytes. Supporting this interpretation, recent studies have reported disadvantage for HLA-E*01:03 individuals in the context of HCMV infection or reactivation after kidney transplantation ^71^, a setting in which NKG2C^+^ lymphocytes may provide critical protection ^7,24,72^. Similarly, HLA-E*01:03 has been associated with greater disease severity in COVID-19 infection ^73,74^.

Increased activation of CD56^dim^ NK cells carrying the NKG2C*02, compared to those with NKG2C*01, in response to HLA-E presenting the LFL peptide, was associated with a stronger downregulation of CD16a. This downregulation represents a functional disadvantage in the context of infection with certain HCMV strains, as it could impair the ADCC potential of NKG2C^+^ NK cells. Notably, our data demonstrate that NKG2C and CD16a exert additive effect on NK cell activation, suggesting that loss of CD16a surface expression could compromise the effectiveness of the adaptive NK response. Furthermore, a recent study reported an association between *NKG2C* CNV and risk of HCMV reactivation in double cord blood transplantation ^20^ supporting a protective role of NKG2C in controlling HCMV in immunocompromised individuals. Given these findings, the clinical impact of *NKG2C* allelic polymorphism should be evaluated in chronical disease, particularly in those involving HCMV reactivation.

Based on the observations from our study, we hypothesize that NKG2C polymorphism and CNV, in conjunction with CD16a and HLA-E polymorphisms, may influence the capacity of aging or immunosuppressed individuals, with impaired adaptive immunity, to control HCMV infection. Furthermore, these genetic factors may contribute to the racial disparities observed in HCMV-related complications following hematopoietic cell transplantation ^75–77^.

## MATERIALS AND METHODS

### Cell Sources and Preparation

Peripheral blood samples were collected from healthy human donors following approval from the Memorial Sloan Kettering Cancer Center (MSKCC) Institutional Review Board (IRB), and donors provided informed, written consent. Buffy coats were obtained from healthy volunteer donors via the New York Blood Center (NYBC, http://nybloodcenter.org/). The MSKCC IRB waived the need for additional research consent for anonymous NYBC samples. Human cytomegalovirus (HCMV) serostatus and self-identified ethnicity and race were provided by NYBC. Peripheral blood mononuclear cells (PBMCs) from each donor were isolated by Ficoll density gradient centrifugation and cryopreserved. Genomic DNA samples were extracted using blood mini kits according to the manufacturer’s instructions (Qiagen); DNA quality and quantity were verified using a NanoDrop Lite Spectrophotometer (ThermoFisher). Human primary fibroblasts cultures were obtained by outgrowth of cells from explanted skin biopsies ^24^. HFs were infected with the HCMV TB40/E strain derivative TB40-BAC_KL7_-SE–enhanced GFP (EGFP) as previously described ^78^. The percentage of infected cells was assessed by flow cytometry (EGFP) before functional experiments.

### KLRC1, KLRC2, KLRK1, KLRD1, HLA-E, HLA-C, FCGR3A and FCGR2A SNP typing

Primers were designed for SSP PCR targeting selected SNPs in various genes (Supplemental Table 1). Each 20μL reaction contained 25-50ng of DNA, 0.5 μL dNTP, 2 μL PCR buffer, and 0.25 μL Taq polymerase (Roche). All reactions were run on Eppendorf Mastercycler Pro-S thermocyclers using the following conditions: 94 °C 2 min, (94°C 15s, 55-63 °C 20s, 72 °C 60-90s) x 40 cycles, 72 °C 1 min. Control primers (either 813 bp or 630 bp) were included in each reaction. Annealing temperature, elongation time, and control primer concentration varied based on primer design (Table1). All PCR products were analyzed using gel electrophoresis on 1.5% agarose gels run for 40 minutes at 125V and imaged with an AlphaImager (AlphaInnotech).

### Phenotypic Analysis by Flow Cytometry

PBMCs (2 × 10^5^ cells per well) were surface stained for 30 min at room temperature with indicated antibodies (Supplemental Table 2) in PBS with 0.5% BSA and 2 mM EDTA for 30 min. Dead cells were excluded by staining with LIVE/DEAD^TM^ (Invitrogen) according to manufacturer instructions, and NK cells were defined by CD3^−^CD56 gated cells. For intracellular staining, the cells were fixed and permeabilized with the FIX & PERM Kit (Invitrogen) according to manufacturer recommendations. For Bcl11b staining’s the ebioscience FOXP3/Transcription factor Staining Kit (Invitrogen) was used according to manufacturer recommendations. All FACS analyses were performed on an LSR Fortessa (BD Biosciences) and analyzed using FlowJo software (10.6.2, Treestar). In all the analyses in this study, MFI refers to mean fluorescence intensity.

### Generation of 721.221 HLA-E and K562 modified cell line

The CML line K562 (ATCC) HLA-ABC deficient BLCL 721.221 (kindly provided by Peter Parham, Stanford University, Stanford, CA) were knocked out for the endogenous HLA-E gene through the LentiCRISPR system; lentivirus was generated as described previously ^24^ using a plasmid encoding an sgRNA targeting HLA-E and Cas9 with puromycin-resistance from Genscript, and viral supernatant was used to transduce the 721.221and K562 cells. An HLA-E-negative clone (EKO) was selected for further use after puromycin selection and single-cell subcloning. Fusion genes containing the leader sequences (amino acids 1-24) of either HLA-A*01 (MA<u>VMAPRTLLL</u>LLSGALALTQTWA) or HLA-G*01 (MV<u>VMAPRTLFL</u>LLSGALTLTETWA) combined with the mature protein of HLA-E were codon-optimized for human translation and synthesized by Genscript. These genes were cloned into the lentivector pERRL ^24^ 5’ of a T2A-eGFP sequence, and the remaining chimeric constructs (A*02:MA<u>VMAPRTLVL</u>LLSGALALTQTWA and C*04:MR<u>VMAPRTLIL</u>LLSGALALTETWA) were derived from the HLA-E:A*01 construct through successive rounds of site-directed mutagenesis (Q5 SDM Kit, NEB). To express the HLA-E chimeras, lentivirus was generated from the pERRL lentivectors and viral supernatant was used to transduce 721.221 EKO; all chimeras were performed simultaneously to avoid any effect of culture time. For functional assays, clones were selected for equal HLA-E surface expression by FACS on a BD Aria followed by single-cell subcloning. All cells were cultured in RPMI 1640 with 10% FCS and Penicillin+Streptomycin.

### NK and T cell Activation Assays

CD107a mobilization and IFN-γ production were used to quantify NK cell activation. Frozen PBMCs were thawed and rested overnight in complete RPMI-1640 medium 10% heat-inactivated fetal bovine serum, 100 U/mL penicillin, 100 μg/mL streptomycin (cRPMI) supplemented with 200 U/mL interleukin-2 (Proleukin, Prometheus Laboratories) and incubated at 37°C with 5% CO_2_. The following day, PBMCs (5 × 10^5^ cells per well) were plated in V-bottom 96 well plates and incubated with target cells at a ratio 5 effectors for 1 Target (5:1) in the presence of anti-CD107a antibody. After 1 hour of co-culture, Brefeldin A (MP Biomedicals) and Golgi stop (BD) were added to each well and incubated for an additional 3 hours.

For P815 cells assays (ATCC), the cells were previously incubated for 20 min at RT with anti CD16a and/or anti-NKG2C mAb, washed and then incubated with PBMC.

For ADCC assays, CD20^+^ Raji cells (ATCC) were incubated with PBMC presence of the anti-CD20 antibody Rituximab (10ng/mL).

For HCMV infected fibroblasts stimulation assays, infected or non-infected fibroblasts were incubated with PBMC at a ratio 5:1 in the presence or absence of HCMV^-^ or HCMV^+^ donor serums (10%).

For plate-bound antibody stimulation assays, 96-well flat-bottom Non-Tissue culture plates were precoated overnight at 4°C in PBS with the following antibodies: anti-NKG2C (134591, R&D Systems), or anti-CD16 (3G8, Biolegend). Wells were washed twice with PBS and Cells (2 × 10^5^) were added to the wells in complete medium containing anti-CD107a antibody. Afterward, cells were washed, and the procedure continued as described above.

For IL-15 stimulation assays, 5-10 million of frozen PBMCs were thawed and incubated in tissue culture treated 6 well plate in cRPMI supplemented with 10 ng/mL rIL-15 and incubated at 37°C with 5% CO_2_ for 64 hours.

### T cell proliferation Assay

Frozen PBMCs were thawed and rested overnight in complete cRPMI supplemented with 100 U/mL IL-2 (Proleukin, Prometheus Laboratories), and incubated at 37°C with 5% CO_2_. The following day, T cells were isolated using the human Pan T isolation kit (Miltenyi) according to the manufacture guidance. T cells were stained with CellTrace Violet (Invitrogen) according to the manufacture guidance. 500 000 T cells with 1 million +/− irradiated 721.221 HL-E KO cells cultured in cRPMI supplemented with IL-15 at 2ng/mL and in absence or presence of anti-CD20 Rituximab (1μg/mL) in a G-Rex 24 Well Plate (Wilsonwoulf) at 8mL/well and incubated for one week at 37°C with 5% CO_2_. At D3, D5 and D7 1mL of co-culture was replaced with fresh media and analyzed by flow cytometry.

### Statistical analysis

Statistical analyses were conducted using unpaired parametric or non-parametric tests to compare independent groups. Before this, a normality test was used to assess whether the groups followed a Gaussian distribution. Depending on the outcome of the normality test, either t-tests or Mann-Whitney tests were used to determine the statistical significance among different groups and mean ± SEM or median ± IQRs are represented. Correlations were assessed using the Pearson correlation coefficient. Comparison of distributions were assessed using Chi-square test. Significance levels are denoted as follows: \**P* ≤ 0.05, \*\**P* ≤ 0.01, *** *P* ≤ 0.001, and \*\*\*\**P* ≤ 0.0001. All analyses and graphical representations were generated using the Prism software (GraphPad).

## LIST OF SUPPLEMENTARY MATERIALS

**Supplemental Table 1.** SSP PCR primers used for genotyping human genomic DNA samples.

**Supplemental Table 2.** Antibodies used for phenotyping and functional analysis.

**Supplemental Fig. 1.** Impact of HCMV infection on lymphocyte population frequencies.

**Supplemental Fig. 2.** Impact of *NKG2C* allelic polymorphism and copy number on HCMV^-/+^ individuals in CD8^+^ T and Vδ1^+^ T cells.

**Supplemental Fig. 3.** NKG2C frequency and MFI stratified by other polymorphism in NK, CD8^+^ T, and Vδ1^+^ T cells.

**Supplemental Fig. 4.** Combine impact of *NKG2C* alleles and copy number, HLA-E variant (rs1264457 G>A), and CD16a variant (rs396991 T>G) on NKG2C^+^ cell frequency in HCMV^+^ individuals.

**Supplemental Fig. 5.** Combined impact of HCMV serostatus and NKG2C allele/copy number on the frequency of NKG2C/A^DP^ CD56^dim^ NK cells.

**Supplemental Fig. 6.** Phenotype and CD107a response of TCRαβ^+^ T cells.

**Supplemental Fig. 7.** CD107a response of NK cell subtypes following activation in a reverse ADCC assay with anti-CD16 loaded P815 target cells.

## ACKNOWLEDGEMENTS

This study was supported by National Institutes of Health (NIH) grants P01 CA23766 and R01 AI125651. Support was also provided by Leukemia & Lymphoma Society, and the NIH P30 CA008748 core grant to Memorial Sloan Kettering Cancer Center.

## AUTHORSHIP CONTRIBUTIONS

Contribution: J-B.L.L. conceived the study. J-B.L.L. and T.K. designed the study. J-B.L.L., T.K., H.L. and R.S. performed experiments. M.K.P provided 721.221 HLA-E modified cell lines. J-B.L.L., T.K., C.M.L. and K.C.H. analyzed and interpreted the data; J-B.L.L., T.K. and K.C.H. wrote and edited the manuscript. All authors reviewed and revised the manuscript and approved the final manuscript.

## REFERENCES

1. Hengel H, Brune W, Koszinowski UH: Immune evasion by cytomegalovirus--survival strategies of a highly adapted opportunist. Trends Microbiol 6:190–7, 1998

2. Pawelec G, Derhovanessian E: Role of CMV in immune senescence. Virus Res 157:175–9, 2011

3. Guma M, Angulo A, Vilches C, et al: Imprint of human cytomegalovirus infection on the NK cell receptor repertoire. Blood 104:3664–71, 2004

4. Lilleri D, Fornara C, Revello MG, et al: Human cytomegalovirus-specific memory CD8+ and CD4+ T cell differentiation after primary infection. J Infect Dis 198:536–43, 2008

5. Boppana SB, Fowler KB: Persistence in the population: epidemiology and transmisson, in Arvin A, Campadelli-Fiume G, Mocarski E, et al (eds): Human Herpesviruses: Biology, Therapy, and Immunoprophylaxis. Cambridge, 2007

6. Hwang I, Zhang T, Scott JM, et al: Identification of human NK cells that are deficient for signaling adaptor FcRgamma and specialized for antibody-dependent immune functions. Int Immunol 24:793–802, 2012

7. Wu Z, Sinzger C, Frascaroli G, et al: Human cytomegalovirus-induced NKG2C(hi) CD57(hi) natural killer cells are effectors dependent on humoral antiviral immunity. J Virol 87:7717–25, 2013

8. Lee J, Zhang T, Hwang I, et al: Epigenetic modification and antibody-dependent expansion of memory-like NK cells in human cytomegalovirus-infected individuals. Immunity 42:431–42, 2015

9. Schlums H, Cichocki F, Tesi B, et al: Cytomegalovirus infection drives adaptive epigenetic diversification of NK cells with altered signaling and effector function. Immunity 42:443–56, 2015

10. Guma M, Budt M, Saez A, et al: Expansion of CD94/NKG2C+ NK cells in response to human cytomegalovirus-infected fibroblasts. Blood 107:3624–31, 2006

11. Rolle A, Pollmann J, Ewen EM, et al: IL-12-producing monocytes and HLA-E control HCMV-driven NKG2C+ NK cell expansion. J Clin Invest 124:5305–16, 2014

12. Perez-Portilla A, Moraru M, Blazquez-Moreno A, et al: Identification of the first cases of complete CD16A deficiency: Association with persistent EBV infection. J Allergy Clin Immunol 145:1288–1292, 2020

13. Beziat V, Liu LL, Malmberg JA, et al: NK cell responses to cytomegalovirus infection lead to stable imprints in the human KIR repertoire and involve activating KIRs. Blood 121:2678–88, 2013

14. Liu LL, Landskron J, Ask EH, et al: Critical Role of CD2 Co-stimulation in Adaptive Natural Killer Cell Responses Revealed in NKG2C-Deficient Humans. Cell Rep 15:1088–1099, 2016

15. Hammer Q, Ruckert T, Borst EM, et al: Peptide-specific recognition of human cytomegalovirus strains controls adaptive natural killer cells. Nat Immunol 19:453–463, 2018

16. Ataya M, Redondo-Pachon D, Llinas-Mallol L, et al: Pretransplant adaptive NKG2C+ NK cells protect against cytomegalovirus infection in kidney transplant recipients. Am J Transplant 20:663–676, 2020

17. Braud VM, Allan DS, O’Callaghan CA, et al: HLA-E binds to natural killer cell receptors CD94/NKG2A, B and C. Nature 391:795–9, 1998

18. Lanier LL, Corliss B, Wu J, et al: Association of DAP12 with activating CD94/NKG2C NK cell receptors. Immunity 8:693–701, 1998

19. Miyashita R, Tsuchiya N, Hikami K, et al: Molecular genetic analyses of human NKG2C (KLRC2) gene deletion. Int Immunol 16:163–8, 2004

20. Cao K, Marin D, Sekine T, et al: Donor NKG2C Copy Number: An Independent Predictor for CMV Reactivation After Double Cord Blood Transplantation. Front Immunol 9:2444, 2018

21. Muntasell A, Lopez-Montanes M, Vera A, et al: NKG2C zygosity influences CD94/NKG2C receptor function and the NK-cell compartment redistribution in response to human cytomegalovirus. Eur J Immunol 43:3268–78, 2013

22. Muntasell A, Pupuleku A, Cisneros E, et al: Relationship of NKG2C Copy Number with the Distribution of Distinct Cytomegalovirus-Induced Adaptive NK Cell Subsets. J Immunol 196:3818–27, 2016

23. Asenjo J, Moraru M, Al-Akioui-Sanz K, et al: NKG2C Sequence Polymorphism Modulates the Expansion of Adaptive NK Cells in Response to Human CMV. HLA 104:e15764, 2024

24. Sottile R, Panjwani MK, Lau CM, et al: Human cytomegalovirus expands a CD8(+) T cell population with loss of BCL11B expression and gain of NK cell identity. Sci Immunol 6:eabe6968, 2021

25. Horowitz A, Guethlein LA, Nemat-Gorgani N, et al: Regulation of Adaptive NK Cells and CD8 T Cells by HLA-C Correlates with Allogeneic Hematopoietic Cell Transplantation and with Cytomegalovirus Reactivation. J Immunol 195:4524–36, 2015

26. Stankovic S, Davey MS, Shaw EM, et al: Cytomegalovirus replication is associated with enrichment of distinct gammadelta T cell subsets following lung transplantation: A novel therapeutic approach? J Heart Lung Transplant 39:1300–1312, 2020

27. Roy Chowdhury R, Valainis JR, Dubey M, et al: NK-like CD8(+) gammadelta T cells are expanded in persistent Mycobacterium tuberculosis infection. Sci Immunol 8:eade3525, 2023

28. Goodier MR, White MJ, Darboe A, et al: Rapid NK cell differentiation in a population with near-universal human cytomegalovirus infection is attenuated by NKG2C deletions. Blood 124:2213–22, 2014

29. Asenjo J, Muntasell A, Lopez-Botet M, et al: Complete genomic characterization of a new KLRC2 allele, NKG2C*03. HLA 98:259–261, 2021

30. Asenjo J, Moraru M, Al-Akioui-Sanz K, et al: Diversity of NKG2C genotypes in a European population: Conserved and recombinant haplotypes in the coding, promoter, and 3’-untranslated regions. HLA, 2022

31. Zhang T, Scott JM, Hwang I, et al: Cutting edge: antibody-dependent memory-like NK cells distinguished by FcRgamma deficiency. J Immunol 190:1402–6, 2013

32. Hatjiharissi E, Xu L, Santos DD, et al: Increased natural killer cell expression of CD16, augmented binding and ADCC activity to rituximab among individuals expressing the Fc{gamma}RIIIa-158 V/V and V/F polymorphism. Blood 110:2561–4, 2007

33. Wu J, Edberg JC, Redecha PB, et al: A novel polymorphism of FcgammaRIIIa (CD16) alters receptor function and predisposes to autoimmune disease. J Clin Invest 100:1059–70, 1997

34. Le Luduec JB, Kontopoulos T, Panjwani MK, et al: Polygenic polymorphism is associated with NKG2A repertoire and influences lymphocyte phenotype and function. Blood Adv 8:5382–5399, 2024

35. Langenbach SY, Zheng L, McWilliams T, et al: Airway vascular changes after lung transplant: potential contribution to the pathophysiology of bronchiolitis obliterans syndrome. J Heart Lung Transplant 24:1550–6, 2005

36. Horowitz A, Djaoud Z, Nemat-Gorgani N, et al: Class I HLA haplotypes form two schools that educate NK cells in different ways. Sci Immunol 1, 2016

37. Poli A, Michel T, Theresine M, et al: CD56bright natural killer (NK) cells: an important NK cell subset. Immunology 126:458–65, 2009

38. van der Ploeg K, Sottile R, Kontopoulos T, et al: Emergence of human CMV-induced NKG2C+ NK cells is associated with CD8+ T-cell recovery after allogeneic HCT. Blood Adv 7:5784–5798, 2023

39. Cocker ATH, Liu F, Djaoud Z, et al: CD56-negative NK cells: Frequency in peripheral blood, expansion during HIV-1 infection, functional capacity, and KIR expression. Front Immunol 13:992723, 2022

40. Cichocki F, Miller JS: Setting traps for NKG2A gives NK cell immunotherapy a fighting chance. J Clin Invest 129:1839–1841, 2019

41. Saez-Borderias A, Romo N, Magri G, et al: IL-12-dependent inducible expression of the CD94/NKG2A inhibitory receptor regulates CD94/NKG2C+ NK cell function. J Immunol 182:829–36, 2009

42. Choi SJ, Koh JY, Rha MS, et al: KIR(+)CD8(+) and NKG2A(+)CD8(+) T cells are distinct innate-like populations in humans. Cell Rep 42:112236, 2023

43. Semmes EC, Nettere DR, Nelson AN, et al: In utero human cytomegalovirus infection expands NK-like FcgammaRIII+CD8+ T cells that mediate Fc antibody functions. J Clin Invest 135, 2024

44. Correia MP, Stojanovic A, Bauer K, et al: Distinct human circulating NKp30(+)FcepsilonRIgamma(+)CD8(+) T cell population exhibiting high natural killer-like antitumor potential. Proc Natl Acad Sci U S A 115:E5980–E5989, 2018

45. Srpan K, Ambrose A, Karampatzakis A, et al: Shedding of CD16 disassembles the NK cell immune synapse and boosts serial engagement of target cells. J Cell Biol 217:3267–3283, 2018

46. Carretero M, Cantoni C, Bellon T, et al: The CD94 and NKG2-A C-type lectins covalently assemble to form a natural killer cell inhibitory receptor for HLA class I molecules. Eur J Immunol 27:563–7, 1997

47. Lanier LL, Corliss BC, Wu J, et al: Immunoreceptor DAP12 bearing a tyrosine-based activation motif is involved in activating NK cells. Nature 391:703–7, 1998

48. Guma M, Busch LK, Salazar-Fontana LI, et al: The CD94/NKG2C killer lectin-like receptor constitutes an alternative activation pathway for a subset of CD8+ T cells. Eur J Immunol 35:2071–80, 2005

49. Fausther-Bovendo H, Wauquier N, Cherfils-Vicini J, et al: NKG2C is a major triggering receptor involved in the V[delta]1 T cell-mediated cytotoxicity against HIV-infected CD4 T cells. AIDS 22:217–26, 2008

50. Yu XX, Shang QN, Liu XF, et al: Donor NKG2C homozygosity contributes to CMV clearance after haploidentical transplantation. JCI Insight 7, 2022

51. Toson B, Michita RT, Matte MCT, et al: Assessment of NKG2C copy number variation in HIV-1 infection susceptibility, and considerations about the potential role of lacking receptors and virus infection. J Hum Genet 67:475–479, 2022

52. Vietzen H, Dohler B, Tran TH, et al: Deletion of the Natural Killer Cell Receptor NKG2C Encoding KLR2C Gene and Kidney Transplant Outcome. Front Immunol 13:829228, 2022

53. Sandor M, Lynch RG: Lymphocyte Fc receptors: the special case of T cells. Immunol Today 14:227–31, 1993

54. Lanier LL, Kipps TJ, Phillips JH: Functional properties of a unique subset of cytotoxic CD3+ T lymphocytes that express Fc receptors for IgG (CD16/Leu-11 antigen). J Exp Med 162:2089–106, 1985

55. Lafont V, Liautard J, Liautard JP, et al: Production of TNF-alpha by human V gamma 9V delta 2 T cells via engagement of Fc gamma RIIIA, the low affinity type 3 receptor for the Fc portion of IgG, expressed upon TCR activation by nonpeptidic antigen. J Immunol 166:7190–9, 2001

56. Groh V, Porcelli S, Fabbi M, et al: Human lymphocytes bearing T cell receptor gamma/delta are phenotypically diverse and evenly distributed throughout the lymphoid system. J Exp Med 169:1277–94, 1989

57. Angelini DF, Borsellino G, Poupot M, et al: FcgammaRIII discriminates between 2 subsets of Vgamma9Vdelta2 effector cells with different responses and activation pathways. Blood 104:1801–7, 2004

58. Bjorkstrom NK, Gonzalez VD, Malmberg KJ, et al: Elevated numbers of Fc gamma RIIIA+ (CD16+) effector CD8 T cells with NK cell-like function in chronic hepatitis C virus infection. J Immunol 181:4219–28, 2008

59. Naluyima P, Lal KG, Costanzo MC, et al: Terminal Effector CD8 T Cells Defined by an IKZF2(+)IL-7R(-) Transcriptional Signature Express FcgammaRIIIA, Expand in HIV Infection, and Mediate Potent HIV-Specific Antibody-Dependent Cellular Cytotoxicity. J Immunol 203:2210–2221, 2019

60. Barbarin A, Cayssials E, Jacomet F, et al: Phenotype of NK-Like CD8(+) T Cells with Innate Features in Humans and Their Relevance in Cancer Diseases. Front Immunol 8:316, 2017

61. Jacquemont L, Tilly G, Yap M, et al: Terminally Differentiated Effector Memory CD8(+) T Cells Identify Kidney Transplant Recipients at High Risk of Graft Failure. J Am Soc Nephrol 31:876–891, 2020

62. Georg P, Astaburuaga-Garcia R, Bonaguro L, et al: Complement activation induces excessive T cell cytotoxicity in severe COVID-19. Cell 185:493–512 e25, 2022

63. Clemenceau B, Vivien R, Debeaupuis E, et al: FcgammaRIIIa (CD16) induction on human T lymphocytes and CD16pos T-lymphocyte amplification. J Immunother 34:542–9, 2011

64. Fauriat C, Ivarsson MA, Ljunggren HG, et al: Education of human natural killer cells by activating killer cell immunoglobulin-like receptors. Blood 115:1166–74, 2010

65. Le Luduec JB, Boudreau JE, Freiberg JC, et al: Novel Approach to Cell Surface Discrimination Between KIR2DL1 Subtypes and KIR2DS1 Identifies Hierarchies in NK Repertoire, Education, and Tolerance. Front Immunol 10:734, 2019

66. Tarek N, Le Luduec JB, Gallagher MM, et al: Unlicensed NK cells target neuroblastoma following anti-GD2 antibody treatment. J Clin Invest 122:3260–70, 2012

67. Kim S, Poursine-Laurent J, Truscott SM, et al: Licensing of natural killer cells by host major histocompatibility complex class I molecules. Nature 436:709–13, 2005

68. Brooks CR, Elliott T, Parham P, et al: The inhibitory receptor NKG2A determines lysis of vaccinia virus-infected autologous targets by NK cells. J Immunol 176:1141–7, 2006

69. Boudreau JE, Hsu KC: Natural killer cell education in human health and disease. Curr Opin Immunol 50:102–111, 2018

70. Bari R, Thapa R, Bao J, et al: KIR2DL2/2DL3-E(35) alleles are functionally stronger than-Q(35) alleles. Sci Rep 6:23689, 2016

71. Guberina H, da Silva Nardi F, Michita RT, et al: Susceptibility of HLA-E*01:03 Allele Carriers to Develop Cytomegalovirus Replication After Living-Donor Kidney Transplantation. J Infect Dis 217:1918–1922, 2018

72. Lopez-Botet M, De Maria A, Muntasell A, et al: Adaptive NK cell response to human cytomegalovirus: Facts and open issues. Semin Immunol 65:101706, 2023

73. Vietzen H, Zoufaly A, Traugott M, et al: Deletion of the NKG2C receptor encoding KLRC2 gene and HLA-E variants are risk factors for severe COVID-19. Genet Med 23:963–967, 2021

74. Hosseini E, Minagar A, Ghasemzadeh M, et al: HLA-E*01:01 + HLA-E*01:01 genotype confers less susceptibility to COVID-19, while HLA-E*01:03 + HLA-E*01:03 genotype is associated with more severe disease. Hum Immunol 84:263–271, 2023

75. Jeurkar C, Leiby B, Banks J, et al: An Examination of Cytomegalovirus, Socioeconomic Status, Race, and Ethnicity on Outcomes after Haploidentical Hematopoietic Transplantation. Transplant Cell Ther 27:327 e1–327 e11, 2021

76. Hamilton BK, Rybicki L, Sekeres M, et al: Racial differences in allogeneic hematopoietic cell transplantation outcomes among African Americans and whites. Bone Marrow Transplant 50:834–9, 2015

77. Boge CLK, Hayes McDonough M, Newman AM, et al: Disparities in Cytomegalovirus Infection Rates by Race and Ethnicity among Pediatric Allogeneic Hematopoietic Cell Transplantation Recipients at a Single Center. Transplant Cell Ther 30:328 e1–328 e12, 2024

78. Sampaio KL, Weyell A, Subramanian N, et al: A TB40/E-derived human cytomegalovirus genome with an intact US-gene region and a self-excisable BAC cassette for immunological research. Biotechniques 63:205–214, 2017

