## Supplementary material for "Polygenic polymorphisms cooperate to drive expansion of HCMV adaptive NK cell and NK-like T cell populations": Supplemetal Material

| SNP | Reaction | Gene region | ARMS primer sequence (Forward & Reverse) | Control primer conc (μM) | Annealing temp (C) | Elongation time (min) |
|---|---|---|---|---|---|---|
| **rs1141715 A>G** | NKG2C V1 | Exon 3 (G) | CTATTACAGTCCTGGAGCAGAACAATCC<br>ATCCACTCCTCAGGACAATGGCCACAGT | 1.25 short | 58 | 1 |
| | NKG2C V2 | Exon 3 (A) | CTATTACAGTCCTGGAGCAGAACAATAT<br>ATCCACTCCTCAGGACAATGGCCACATT | 1.25 short | 58 | 1 |
| **rs28403159 A>G** | NKG2C V3 | Exon 1 (C) | GTATTTCTAAATACAGTATTTTGATTGTTGAC<br>CTCACTTCTGAGAAGGTTCCTCTTTGTTTTC | 0.833 long | 50 | 1 |
| | NKG2C V4 | Exon 1 (T) | GTATTTCTAAATACAGTATTTTGATTGTTGAC<br>CTCACTTCTGAGAAGGTTCCTCTTTGTTTTT | 0.833 long | 50 | 1 |
| **N/A** | NKG2C Neg | | GTAGGATATACTTCGGATTTCTATTTGATGC<br>GCTAAAATTAAATACAAGTGATGTATAAGAAAAAG | 1.25 long | 58 | 1 |
| **rs2246809 A>G** | NKG2D V1 | 5' UTR (A) | GCCTGTTTAAATTTATTAATGCACAGGAGATTCT<br>GCAATACAACAAACAGAAATCTGAGTAACCTC | 5 long | 58 | 1 |
| | NKG2D V2 | 5' UTR (G) | GCCTGTTTAAATTTATTAATGCACAGGAGATTCC<br>GCAATACAACAAACAGAAATCTGAGTAACCTC | 5 long | 58 | 1 |
| **rs11053781 C>T** | NKG2D V3 | Intron 2 (C) | CATAAAATCTTTGCCTAGACTAATGTCCGG<br>ATCACGTTACCTGACTTCAATATATACGA | 1.25 long | 57 | 1 |
| | NKG2D V4 | Intron 2 (T) | CATAAAATCTTTGCCTAGACTAATGTCCAA<br>ATCACGTTACCTGACTTCAATATATACGA | 1.25 long | 57 | 1 |
| **rs1049174 G>C** | NKG2D V5 | 3' UTR (G) | AGTGAAGGAAGAGAAGGCCAGCAGATCTC<br>ACAAGTACAAATCATAGCAAAGGAAACGTCCTAAGATC | 2.5 long | 60 | 1 |
| | NKG2D V6 | 3' UTR (C) | AGTGAAGGAAGAGAAGGCCAGCAGATCCG<br>ACAAGTACAAATCATAGCAAAGGAAACGTCCTAAGATC | 2.5 long | 60 | 1 |
| **rs2302489 A>T** | CD94 V1 | Exon (A) | GAACATCATTTAAATACACAATTTTTCATTCTCAA<br>CAATGAAGAATCAGGAAAATATAATGAATGTGG | 1.25 long | 56 | 1 |
| | CD94 V2 | Exon (T) | GAACATCATTTAAATACACAATTTTTCATTCTCCT<br>CAATGAAGAATCAGGAAAATATAATGAATGTGG | 2.5 long | 56 | 1 |
| **rs396991 T>G** | CD16 V1 low affinity | Exon 4 (T) | AGCGGCTCCTACTTCTGCAGGGGGCTCT<br>CAGAAAGAGACTTGGTACCCAGGTGGTA | 1.66 long | 63 | 1.5 |
| | CD16 V2 high affinity | Exon 4 (G) | AGCGGCTCCTACTTCTGCAGGGGGCTCG<br>CAGAAAGAGACTTGGTACCCAGGTGGTA | 1.66 long | 63 | 1.5 |
| **rs1801274 A>G** | CD32 V1 high affinity | Exon 4 (A) | CAGAATGGAAAATCCCAGAAATTCTCCGA<br>ACAGTGATGGTCACAGGCTTGGATGATA | 2.5 long | 63 | 1 |
| | CD32 V2 low affinity | Exon 4 (G) | CAGAATGGAAAATCCCAGAAATTCTCCGG<br>ACAGTGATGGTCACAGGCTTGGATGATA | 2.5 long | 63 | 1 |
| **rs11586238 C>G** | CD2 V1 | Upstream(C) | GTGTTCTGCAGGGATCTCTGAGGCTCA<br>TTTTGCTTGGTTCCCATTGCCTTCAGCATCC | 5 long | 63 | 1 |

| | | | | | | |
|---|---|---|---|---|---|---|
| | CD2 V2 | Upstream(G) | GTGTTCTGCAGGGATCTCTGAGGCTCA<br>TTTTGCTTGGTTCCCATTGCCTTCAGCATCG | 5 long | 63 | 1 |
| **rs12044852 C>A** | CD58 V1 | Intron 1 (C) | GCTTGAGGATACACACGTGATTCCTAACAAC<br>CAAACACCCTGCTCTGGTCTGTCCTGT | 2.5 long | 63 | 1 |
| | CD58 V2 | Intron 1 (A) | GCTTGAGGATACACACGTGATTCCTAACAAA<br>CAAACACCCTGCTCTGGTCTGTCCTGT | 2.5 long | 63 | 1 |
| **rs1264457 G>A** | HLA-E 01:01 | Exon 2 (A) | ATGCATGGCTGCGAGCTGGGGCCCGATA<br>GCATGTGTCTTCCAGGTAGGCTCTCTCG | 10 long | 63 | 1 |
| | HLA-E 01:03 | Exon 2 (G) | ATGCATGGCTGCGAGCTGGGGCCCGATG<br>GCATGTGTCTTCCAGGTAGGCTCTCTCG | 10 long | 63 | 1 |
| **rs17408553 G>T** | HLA-C1 | Exon 2 (G) | TTGGGAGGGAAACGGCCTCTGCGAA<br>TTGGGAGGGAAACGGCCTCTGGGAA<br>GCTCTGGTTGTAGTAGCCGCGCATG | 10 long | 63 | 1 |
| | HLA-C2 | Exon 2 (T) | TTGGGAGGGAAACGGCCTCTGCGAA<br>TTGGGAGGGAAACGGCCTCTGGGAA<br>GCTCTGGTTGTAGTAGCCGCGCAAT | 10 long | 63 | 1 |
| **N/A** | Long control | | CCAAGCCCAACCTTAAGAAGAAAATTGGAG<br>CCAAACCCACGGTACGCATGGGAACACTGC | - | | |
| **N/A** | Short control | | CCCACCTTCCCCTCTCTCCAGGCAAATGGG<br>GGGCCTCAGTCCCAACATGGCTAAGAGGTG | - | | |

**Supplemental Table 1.** SSP PCR primers used for genotyping human genomic DNA samples. Each is listed with SNP, its associated reaction name or variant (referred to throughout this study), targeted gene region, primer sequences, control primer concentration, annealing temperature, and elongation time.

| Reagent | Color | Clone | Company | Isotype |
|---|---|---|---|---|
| Dapi | Dapi | | BD | |
| CD3 | BV650 | UCHT1 | BD | IgG1 Mouse, K |
| CD56 | ECD | N901 | IOTest (BC) | IgG1 Mouse, K |
| NKG2C | APC/FITC | REA205 | Miltenyi | REA |
| NKG2C | Unconjugated | 134591 | R&D | IgG Mouse |
| NKG2A | PeCy7 | Z199 | IOTest (BC) | IgG2b Mouse, K |
| KIR2DL1/S1 | PeCy5.5 | EB6B | IOTest (BC) | IgG1 Mouse, K |
| KIR2DL2/L3/S2 | BV605 | CH-L | BD | IgG2b Mouse, K |
| KIR3DL1 | BV711/APC | DX9 | BD | IgG1 Mouse, K |
| KIR3DL1/S1 | APCVio770 | REA168 | Miltenyi | REA |
| CD107a | BV786/BV421 | H4A3 | BD | IgG1 Mouse, K |
| IFN-$\gamma$ | AF700 | B27 | BD | IgG1 Mouse, K |
| CD94 | APC | NA | Miltenyi | REA |
| TCR αβ | BV711 | IP26 | Biolegend | IgG1 Mouse, K |
| TCR γδ | PE | B1 | Biolegend | IgG1 Mouse, K |
| CD4 | Alexa Fluor 700 | RPA-T4 | Biolegend | IgG1 Mouse, K |
| CD8a | BV786/BV570 | RPA-T8 | BD | IgG1 Mouse, K |
| Vd1 | APCVio770 | REA173 | Miltenyi | REA |
| Vd2 | PerCP-Vio 700 | REA771 | Miltenyi | REA |
| CD16 | BV786/BV421/Unconjugated | 3G8 | Biolegend | IgG1 Mouse, K |
| FcεRIγ | FITC | NA | Millipore | IgG Rabbit |
| Bcl11b | FITC | 25B6 | abcam | IgG2A Rat, K |
| Isotype Bcl11b | FITC | Ab18446 | abcam | IgG2A Rat, K |

**Supplemental Table 2.** Antibodies used for phenotyping and functional analysis.

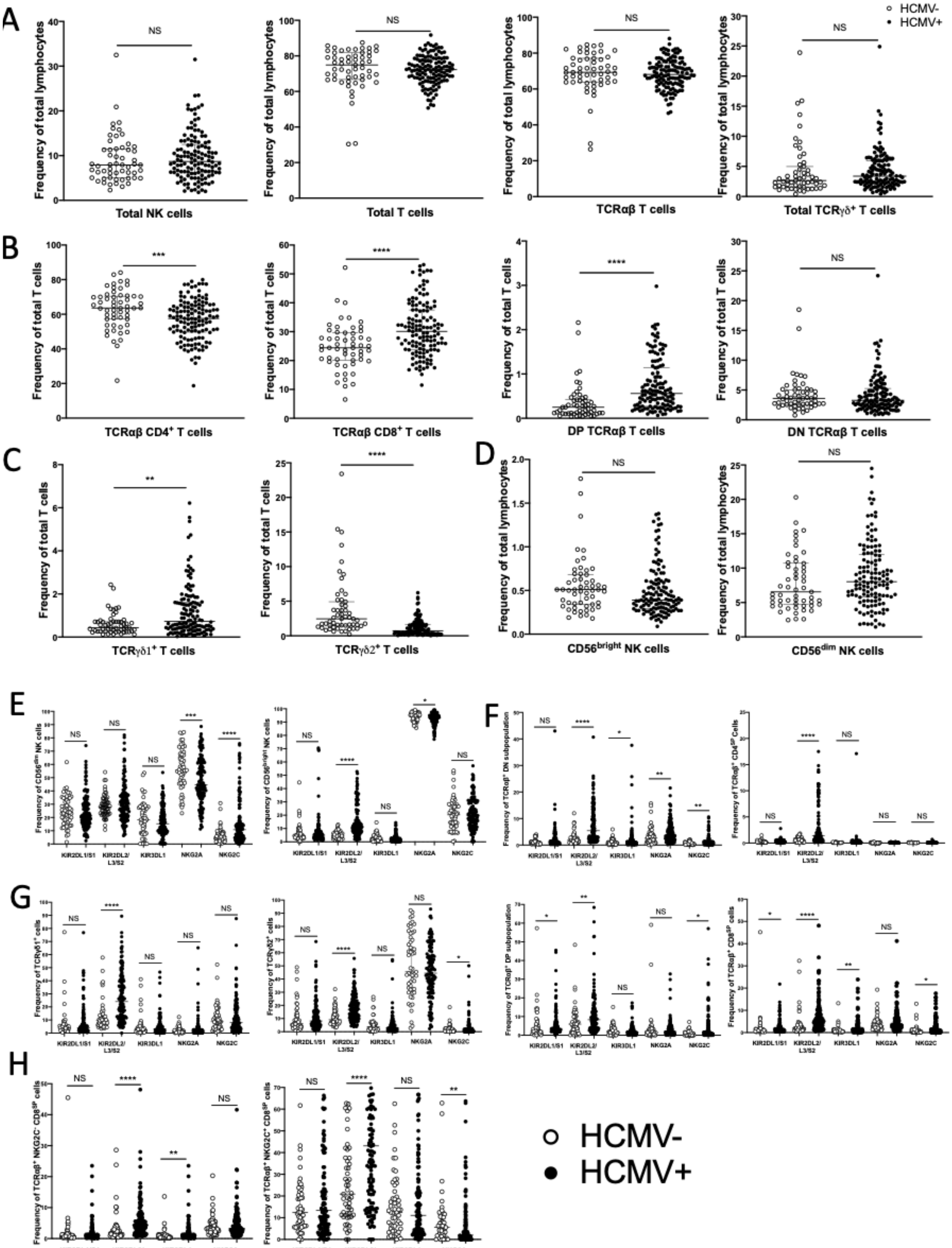


**Supplemental Fig. 1. Impact of HCMV infection on lymphocyte population frequencies.** (A)

Frequency of lymphocytes populations stratified by HCMV serostatus. (B) Frequency of TCRαβ$^+$ T cell subsets stratified by HCMV serostatus. (C) Frequency of TCRγδ$^+$ T cell subsets stratified by HCMV serostatus. (D) Frequency of NK cell subsets stratified by HCMV serostatus. (E-G) KIR2DL1/S1, KIR2DL2/L3/S2, KIR3DL1, NKG2A and NKG2C frequency on indicated lymphocyte subsets stratified by HCMV serostatus. Mann-Whitney tests were performed and median ± IQRs presented in panels A-D. Symbols represent individual samples. *$p < 0.05$, **$p < 0.01$, ***$p < 0.001$, and ****$p < 0.0001$.

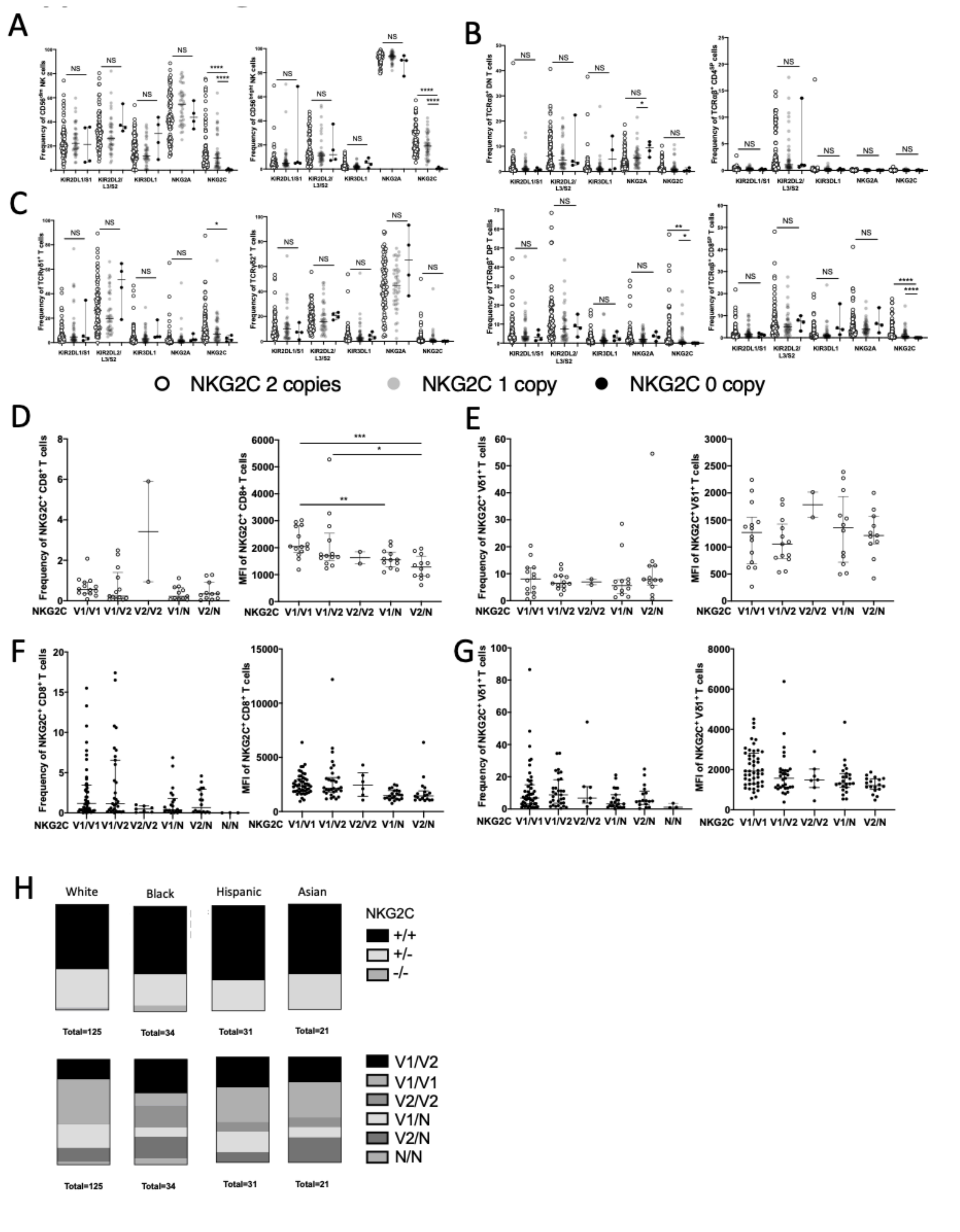


**Supplemental Fig. 2. Impact of *NKG2C* allelic polymorphism and copy number on HCMV$^{-/+}$ individuals in CD8$^{+}$ T and Vδ1$^{+}$ T cells.** (A-C) KIR2DL1/S1, KIR2DL2/L3/S2, KIR3DL1,

NKG2A and NKG2C frequency on indicated lymphocyte subsets stratified NKG2C copy number. (D) NKG2C frequency and MFI stratified by *NKG2C* variant and copy number in $CD8^+$ T cells from $HCMV^-$ individuals. (E) NKG2C frequency and MFI stratified by *NKG2C* variant and copy number in $V\delta1^+$ T cells from $HCMV^-$ individuals. (F) NKG2C frequency and MFI stratified by *NKG2C* variant and copy number in $CD8^+$ T cells from $HCMV^+$ individuals. (G) NKG2C frequency and MFI stratified by *NKG2C* variant and copy number in $V\delta1^+$ T cells from $HCMV^+$ individuals. (H) Distribution of *NKG2C* copy number and rs1141715 alleles among different ethnic groups in our cohort. Mann-Whitney tests were performed and median ± IQRs presented in panels A-G. Symbols represent individual samples. $^*p < 0.05$, $^{**}p < 0.01$, $^{***}p < 0.001$, and $^{****}p < 0.0001$.

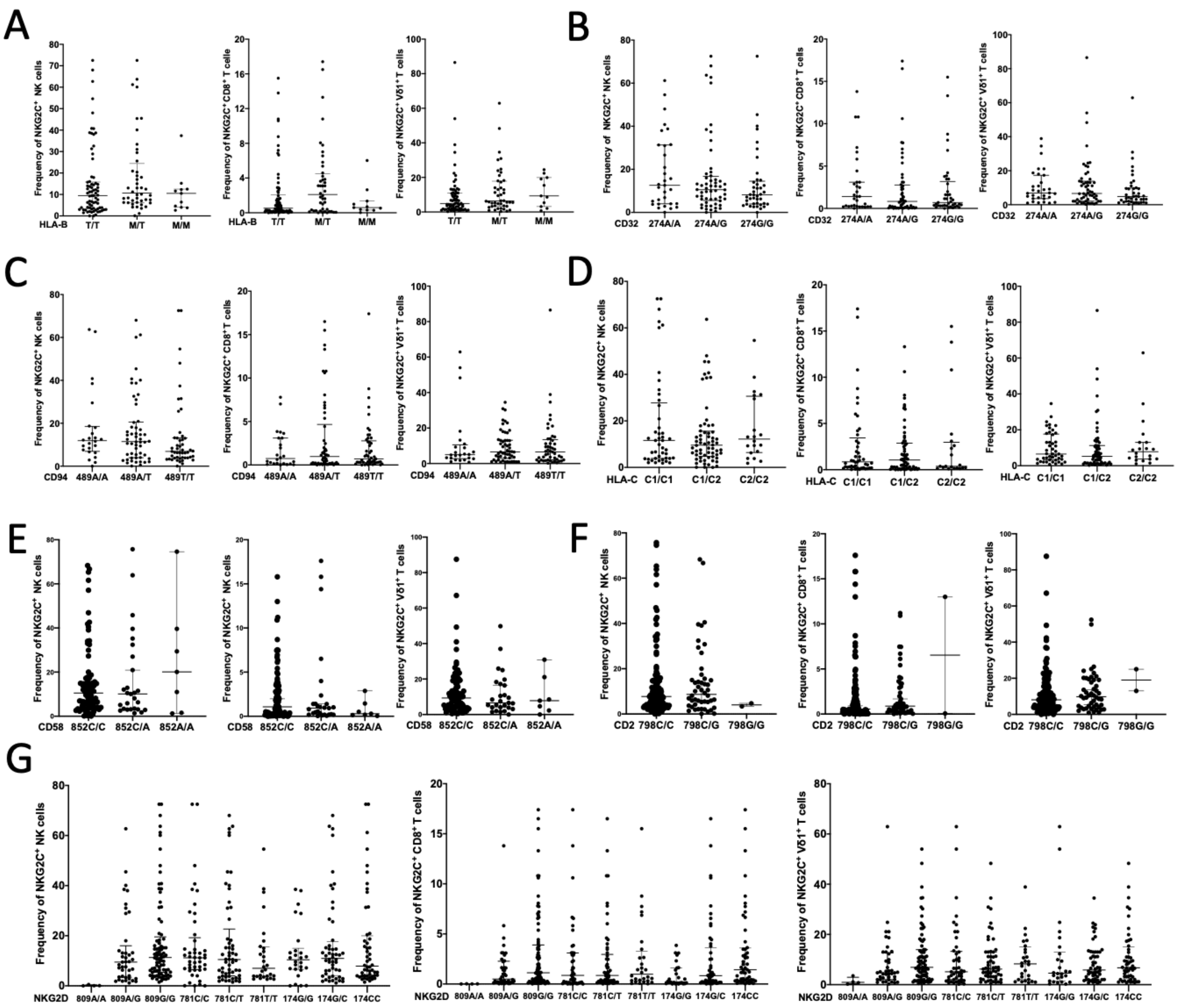


**Supplemental Fig. 3. NKG2C frequency and MFI stratified by other polymorphism in NK, CD8⁺ T, and Vδ1⁺ T cells**. NKG2C frequency NK, CD8⁺ T, and Vδ1⁺ T cells from HCMV⁺ individuals stratified by: (A) HLA-B -21 variants, (B) CD32a polymorphism (rs1801274 A>G), (C) CD94 polymorphism (rs3759272 G>T), (D) HLA-C polymorphism (rs17408553 G>T), (E) CD2 polymorphism (rs11586238 C>G), (F) CD58 polymorphism (rs12044852 C>A), (G) NKG2D polymorphism (rs2246809 A>G, rs11053781 C>T, and rs1049174 G>C. Mann-Whitney tests were performed and median ± IQRs presented in panels A-E. Symbols represent individual samples. **p* < 0.05, ***p* < 0.01, ****p* < 0.001, and *****p* < 0.0001.

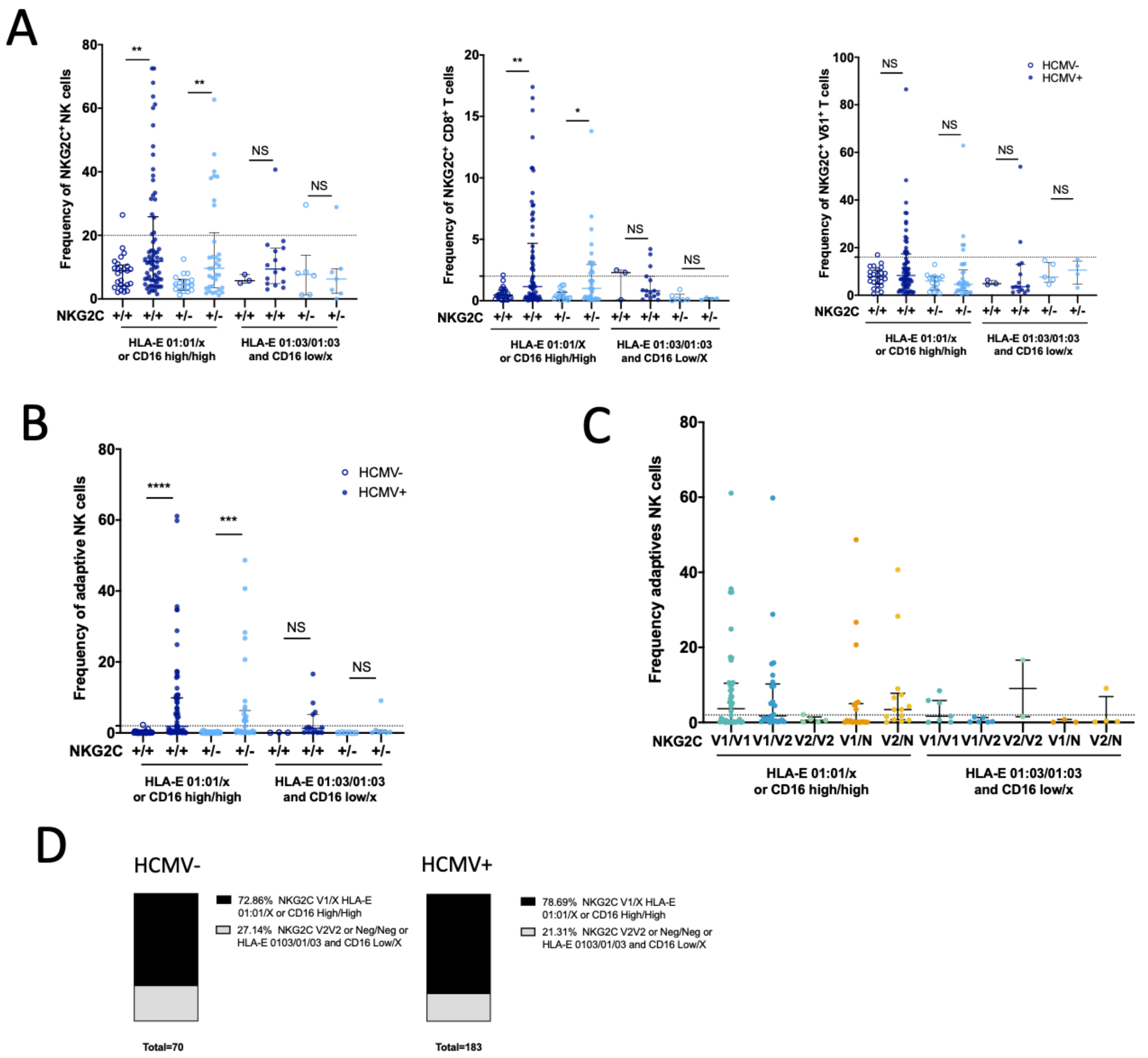


**Supplemental Fig. 4. Combine impact of *NKG2C* alleles and copy number, HLA-E variant (rs1264457 G>A), and CD16a variant (rs396991 T>G) on NKG2C$^+$ cell frequency in HCMV$^+$ individuals.** (A) NKG2C frequency stratified by the combine polymorphism of *NKG2C* copy number HLA-E variants (rs1264457 G>A) and CD16a variants (rs396991 T>G) in NK cells, TCRαβ$^+$CD8$^+$ T cells and Vδ1$^+$ T cells from HCMV$^-$ and HCMV$^+$ individuals. (B) Combine impact of *NKG2C* copy number, HLA-E variant, and CD16a variant on NKG2C$^+$ FcεRIγ$^-$ adaptive NK cell frequency in HCMV$^-$ and HCMV$^+$ individuals. (C) Combine impact of *NKG2C* copy number and allelic variant, HLA-E variant, and CD16a variant on NKG2C$^+$ FcεRIγ$^-$ adaptive NK cell frequency in $HCMV^-$ and $HCMV^+$ individuals. (D) Distribution of the combine association of *NKG2C* alleles, CD16a and HLA-E alleles among $HCMV^-$ and $HCMV^+$ individuals in our cohort. Mann-Whitney tests were performed and median ± IQRs presented in panels A and B. Comparison of distributions were assessed using Chi-square test in panel C. Symbols represent individual samples. *$p < 0.05$, **$p < 0.01$, ***$p < 0.001$, and ****$p < 0.0001$.

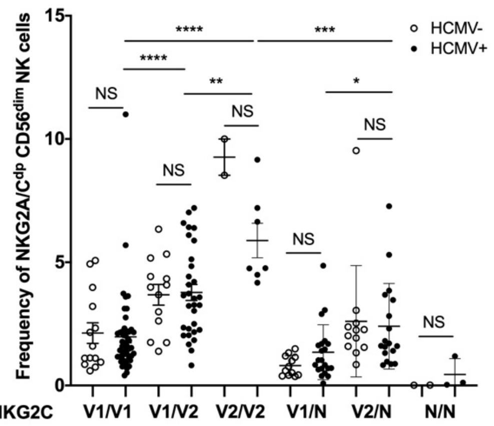


**Supplemental Fig. 5. Combined impact of HCMV serostatus and NKG2C allele/copy number on the frequency of NKG2C/A$^{DP}$ CD56$^{dim}$ NK cells.** Mann-Whitney tests were performed and median ± IQRs presented. Symbols represent individual samples. *$p < 0.05$, **$p < 0.01$, ***$p < 0.001$, and ****$p < 0.0001$.

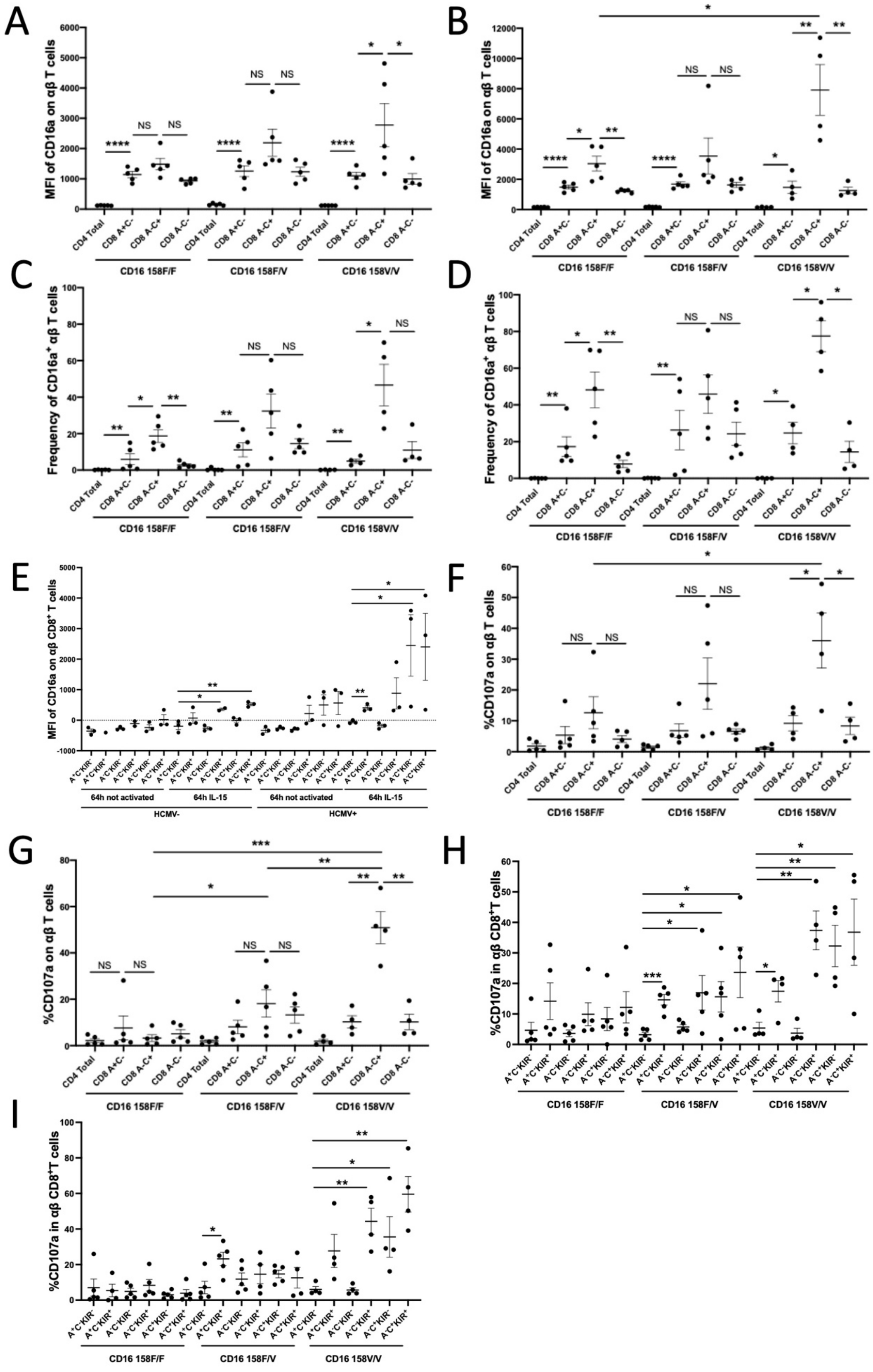
A
MFI of CD16a on αβ T cells
CD4 Total
CD8 A+C-
CD8 A-C+
CD8 A-C-
CD16 158F/F
CD16 158F/V
CD16 158V/V
B
MFI of CD16a on αβ T cells
C
Frequency of CD16a⁺ αβ T cells
D
Frequency of CD16a⁺ αβ T cells
E
MFI of CD16a on αβ CD8⁺ T cells
64h not activated
64h IL-15
HCMV-
HCMV+
F
%CD107a on αβ T cells
G
%CD107a on αβ T cells
H
%CD107a in αβ CD8⁺T cells
I
%CD107a in αβ CD8⁺T cells
NS

**Supplemental Fig. 6. Phenotype and CD107a response of TCRαβ$^+$ T cells**. (A) CD16a MFI of CD4$^+$ and CD8$^+$ T cells subpopulations from HCMV$^+$ individuals, grouped by CD16a polymorphisms, defined by NKG2A and NKG2C positivity, before IL-15 stimulation. (B) CD16a MFI of TCRαβ$^+$ CD4$^+$ and CD8$^+$ T cells subpopulation from HCMV$^+$ individuals, grouped by CD16a polymorphisms, defined by NKG2A and NKG2C positivity, after 64 hours of IL-15 stimulation. (C) Frequency of CD16a$^+$ CD4$^+$ and CD8$^+$ T cells subpopulation from HCMV$^+$ individuals, grouped by CD16a polymorphisms, defined by NKG2A and NKG2C positivity, before IL-15 stimulation. (D) Frequency of CD16a$^+$ CD4$^+$ and CD8$^+$ T cells subpopulation from HCMV$^+$ individuals, grouped by CD16a polymorphisms, defined by NKG2A and NKG2C positivity, after 64 hours of IL-15. (E) CD16a MFI of CD8$^+$ T cells subpopulation from HCMV$^-$ or HCMV$^+$ CD16a 158V/V individuals, defined by NKG2A, NKG2C and KIR positivity, before and after 64 hours of IL-15 stimulation. (F) CD107a response of CD4$^+$ or CD8$^+$ T cells subpopulation from HCMV$^+$ individuals with different CD16a polymorphisms, defined by NKG2Aand NKG2C positivity, after 64 hours of IL-15 stimulation, co-cultured with anti-CD16a and P815 cells for 4 hours. (G) CD107a response of CD4$^+$ or CD8$^+$ T cells subpopulation from HCMV$^+$ individuals with different CD16a polymorphisms, defined by NKG2A and NKG2C positivity, after 64 hours of IL-15 stimulation, co-cultured with Raji cells and Rituximab mAb. (H) CD107a response of CD8$^+$ T cells subpopulation from HCMV$^+$ individuals with different CD16a polymorphisms, defined by NKG2A, NKG2C and KIR positivity, after 64 hours of IL-15 stimulation, co-cultured with anti-CD16a and P815 cells for 4 hours. (I) CD107a response of CD8$^+$ T cells subpopulation from HCMV$^+$ individuals with different CD16a polymorphisms, defined by NKG2A, NKG2C and KIR positivity, after 64 hours of IL-15 stimulation, co-cultured with Raji cells and Rituximab. Mann-

Whitney tests were performed and median ± IQRs presented in panels A-F. Symbols represent individual samples. *$p < 0.05$, **$p < 0.01$, ***$p < 0.001$, and ****$p < 0.0001$.

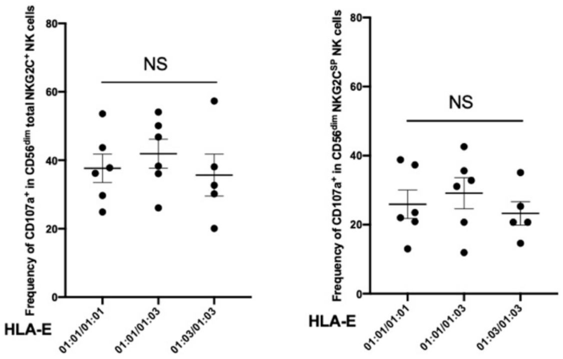


**Supplemental Fig. 7. CD107a response of NK cell subtypes following activation in a reverse ADCC assay with anti-CD16 loaded P815 target cells.** T-tests were performed and mean ± SEM are presented. Symbols represent individual samples. *$p < 0.05$, **$p < 0.01$, ***$p < 0.001$, and ****$p < 0.0001$.
